# Contrasting parental roles shape sex differences in poison frog space use but not navigational performance

**DOI:** 10.1101/2022.05.21.492915

**Authors:** Andrius Pašukonis, Shirley Jennifer Serrano-Rojas, Marie-Therese Fischer, Matthias-Claudio Loretto, Daniel A. Shaykevich, Bibiana Rojas, Max Ringler, Alexandre-Benoit Roland, Alejandro Marcillo-Lara, Eva Ringler, Camilo Rodríguez, Luis A. Coloma, Lauren A. O’Connell

## Abstract

Sex differences in vertebrate spatial abilities are typically interpreted under the adaptive specialization hypothesis, which posits that male reproductive success is linked to larger home ranges and better navigational skills. The androgen spillover hypothesis counters that enhanced male spatial performance may be a byproduct of higher androgen levels. Animal groups that include species where females are expected to outperform males based on life-history traits are key for disentangling these hypotheses. We investigated the association between sex differences in reproductive strategies, spatial behavior, and androgen levels in three species of poison frogs. We tracked individuals in natural environments to show that contrasting parental sex roles shape sex differences in space use, where the sex performing parental duties shows wider-ranging movements. We then translocated frogs from their home areas to test their navigational performance and found that the caring sex outperformed the non-caring sex only in one out of three species. In addition, males across species displayed more explorative behavior than females. Furthermore, androgen levels correlated with explorative behavior and homing accuracy. Our findings suggest that poison frog reproductive strategies shape space use patterns but not navigational performance, providing counterevidence to the prevailing view of adaptive sex differences in spatial abilities.

## Résumé (français)

Chez les vertébrés, les différences de capacités spatiales entre les sexes des sont généralement interprétées selon l’hypothèse de la “spécialisation adaptative”, selon laquelle le succès reproductif des mâles serait lié à l’utilisation d’un territoire plus étendu leur procurant de meilleures aptitudes à la navigation. Cependant, une hypothèse du “débordement androgénique” propose que l’amélioration des performances spatiales des mâles puisse être en fait, un sous-produit de niveaux d’androgènes plus élevés. Pour différencier ces hypothèses, il semble donc nécessaire d’utiliser des groupes d’animaux pour lesquels il existe des espèces dont les femelles seraient plus performantes que les mâles en raison de leur écologie. Ainsi nous avons étudié les différences entre les sexes et leur interaction avec les stratégies de reproduction, le comportement spatial et les niveaux d’androgènes, chez trois espèces de grenouilles dendrobatoidés *sensu lato*. Nous avons donc suivi des individus dans leurs environnements naturels, et montré que les rôles parentaux déterminent les différences sexuelles dans l’utilisation de l’espace, le sexe en charge des fonctions parentales se déplaçant sur de plus grandes distances. Nous avons ensuite déplacé les grenouilles de leur site d’origine afin de tester leurs performances navigationnelles et nous avons constaté que le sexe en charge des soins parentaux surpasse le sexe qui ne prodigue pas de soins chez une seule des trois espèces. De plus, chez les trois espèces, les mâles ont montré un comportement exploratoire supérieur à celui des femelles. Enfin, les niveaux d’androgènes sont corrélés au comportement exploratoire et à la précision de la trajectoire. Nos résultats suggèrent donc que les stratégies de reproduction des grenouilles dendrobatoidés influencent l’utilisation de l’espace mais pas les performances de navigation, en contraction avec l’hypothèse dominante sur les différences adaptatives entre les sexes dans les capacités spatiales.

## Resumen (español)

Las diferencias en las habilidades espaciales de machos y hembras en especies de vertebrados se han interpretado comúnmente bajo la hipótesis de la “especialización adaptativa”, la cual sugiere que el éxito reproductivo de los machos está ligado a ámbitos hogareños más extensos y a mejores habilidades de navegación. Por otra parte, la hipótesis de la “sobreproducción de andrógenos” propone que un mejor desempeño espacial en los machos podría ser un subproducto de niveles de andrógenos más altos. Estudios que involucren grupos de animales con especies en las que se espera que las hembras superen el desempeño de los machos con base en ciertos rasgos de la historia de vida son cruciales para poder discernir entre estas dos hipótesis. En este estudio investigamos la asociación entre las diferencias en estrategias reproductivas de machos y hembras, el comportamiento espacial, y los niveles de andrógenos en tres especies de ranas venenosas. Seguimos individuos en su ambiente natural para mostrar que los roles contrastantes de machos y hembras en cuanto al comportamiento de cuidado parental moldean las diferencias en uso del espacio entre los sexos, de forma que el sexo encargado de llevar a cabo el cuidado parental presenta movimiento en un rango mucho más amplio. Luego, translocamos ranas a lugares fuera de su ámbito hogareño para investigar su desempeño de navegación y encontramos que el sexo encargado del cuidado parental superó el desempeño del sexo opuesto únicamente en una de las tres especies estudiadas. Adicionalmente, los machos de las tres especies tuvieron un comportamiento exploratorio más marcado que las hembras. Asimismo, encontramos que los niveles de andrógenos se correlacionan con el comportamiento exploratorio y con la precisión para regresar al hogar. Nuestros hallazgos sugieren que las estrategias reproductivas de las ranas venenosas moldean los patrones de uso del espacio pero no el desempeño de navegación, proporcionando

## Introduction

Sex differences in spatial abilities are well established in mammals, where males tend to have larger home ranges and enhanced navigational skill compared to females (Clint et al., 2012; Gray & Buffery, 1971; Jonasson, 2005; Jones et al., 2003). In a series of comparative studies, sex differences in space use have been linked to reproductive strategy, where polygamous rodents show sex differences in home range size and spatial abilities, but monogamous species do not (Galea et al., 1994; Gaulin et al., 1990; Gaulin & FitzGerald, 1986; Gaulin & Fitzgerald, 1989; Sawrey et al., 1994). Furthermore, across many human ethnic groups, men tend to score higher on spatial tests related to 3D mental rotations, whereas women tend to score better on object location memory (Eals & Silverman, 1994; Silverman et al., 2007; reviewed in Clint et al., 2012; Jones et al., 2003). The adaptive specialization hypothesis has been widely used to interpret these sex differences by arguing that enhanced spatial abilities in males are an adaptive trait linked to fitness, where males with better navigational skills and larger home ranges may have increased reproductive success (Gaulin & FitzGerald, 1986; Gaulin & Fitzgerald, 1989; Jones et al., 2003). In addition, maternal care in mammals may limit space use and exploration in females (Barnett & McEwan, 1973; Sherry & Hampson, 1997; Trivers, 1972). With few exceptions (Costa et al., 2011; Guigueno et al., 2014; Perry & Garland Jr, 2002; Sherry et al., 1993), empirical support for adaptive sex differences in spatial abilities is based on research in mammals, where males typically have larger home ranges than females (but see Mabry et al., 2013; Mysterud et al., 2001; Ofstad et al., 2016). Taxonomically diverse study systems are needed to test the adaptive specialization hypothesis and its broader implications for the evolution of vertebrate spatial cognition.

Clint et al. (2012) challenged the widely accepted adaptive explanations of sex differences. They countered that sex differences in spatial behavior might be a byproduct of sex differences in androgens rather than an adaptation based on reproductive strategies (i.e., the androgen spill-over hypothesis). Higher androgen levels in mammals enhance spatial performance through effects on neural development and plasticity (Dawson et al., 1975; Galea et al., 1995; Isgor & Sengelaub, 1998; Joseph et al., 1978; Neave et al., 1999; Roof & Havens, 1992; Schulz & Korz, 2010; Sherry & Hampson, 1997; Stewart et al., 1975; van Goozen et al., 1995; Williams et al., 1990). In humans, female performance in spatial ability tasks correlates positively with androgen levels and improves with androgen treatments (Aleman et al., 2004; Burkitt et al., 2007; Driscoll et al., 2005). From an adaptationist perspective, this relationship has been viewed as the proximate mechanism for the selection on males’ spatial abilities. However, Clint et al. (2012) argued that better spatial abilities in males might be a byproduct of sex differences in androgens unrelated to selective pressures on spatial behavior and reproductive success. As mammals have limited diversity in reproductive strategies, we lack comparisons with species where females, which have lower androgen levels, have larger home ranges and are expected to have better spatial abilities than males. To disentangle the effect of androgens and life-history traits on sex differences in spatial behavior, we need comparative research in groups of animals where either males or females among closely related species have more complex spatial behavior.

Fish and amphibians show a remarkable variety of mating strategies and parental sex roles compared to mammals and birds, including widespread polyandry and male uniparental care (Duellman, 1989; Gross & Sargent, 1985; Helfman et al., 2009; Schulte et al., 2020; Wells, 1977; Zamudio et al., 2016). Such behavioral diversity provides natural comparison groups to test alternative hypotheses about sex differences in spatial abilities. In Neotropical poison frogs (Dendrobatoidea), male and female uniparental care, biparental care, and flexible parental sex roles occur among closely related species (Schulte et al., 2020; Summers & Tumulty, 2014; Weygoldt, 1987). Poison frog parental care involves complex spatial behavior, where parents navigate the rainforest to transport tadpoles from terrestrial clutches to pools of water (Beck et al., 2017; Pašukonis et al., 2019; E. Ringler et al., 2013). Poison frogs show well-developed spatial cognition and rely on spatial memory to relocate home territories and tadpole deposition sites (Beck et al., 2017; Liu et al., 2016, 2019; Pašukonis et al., 2014, 2016; Stynoski, 2009). Male tadpole transport is the ancestral and most common form of parental care in poison frogs (Carvajal-Castro et al., 2021; Summers & Tumulty, 2014; Weygoldt, 1987), but female transport and flexible parental roles have evolved in some species (E. K. Fischer & O’Connell, 2020; E. Ringler, Pašukonis, et al., 2015). In species where females perform parental care, frogs place their tadpoles in small, resource-poor nurseries. Females frequently return to these pools to supplement the tadpoles’ diet by provisioning with unfertilized eggs (Brust, 1990; Summers & Tumulty, 2014). All these parental behaviors require well-developed spatial memory and navigation abilities. Therefore individuals with better spatial memory may have increased reproductive fitness, leading to enhanced spatial abilities in the sex that performs parental care, as proposed by the adaptive specialization hypothesis. Poison frogs also show sex-typical differences in androgen levels, where males have higher androgen levels than females, although androgen levels decrease during tadpole transport in males (E. K. Fischer & O’Connell, 2020). Androgens are also elevated in response to territorial intrusions in males, suggesting that androgens are regulated in these amphibians similarly to other vertebrate taxa (Rodríguez et al., 2022). Thus, comparative studies in poison frogs provide a unique opportunity to understand how parental roles and reproductive strategies shape sex differences in space use and navigational abilities, and how hormones regulate these behaviors.

Here we report extensive field studies on sex differences in spatial behavior across three poison frog species with contrasting parental sex roles and reproductive strategies: the Brilliant-Thighed Poison Frog *Allobates femoralis*, an inconspicuous frog with flexible but predominantly male parental care, the Dyeing Poison Frog *Dendrobates tinctorius*, an aposematic frog with obligate male care, and the Diablito Poison Frog *Oophaga sylvatica*, an aposematic species with obligate female care. We tracked frogs with miniature tags to quantify sex differences in home range and parental care-associated space in their natural environment. We then quantified the sex differences in navigational performance by experimentally translocating frogs from their home areas and tracking their homing behavior. We also used noninvasive methods to measure androgen levels before and after translocation. Based on the adaptive specialization hypothesis, we predicted that the tadpole transporting sex (males in *D. tinctorius* and *A. femoralis*, females in *O. sylvatica*) would have wider-ranging space use and better navigational performance. Following the androgen spillover hypothesis, we predicted that males would show enhanced navigation regardless of species differences in reproductive strategy.

## Results

### Sex differences in parental roles predict sex differences in space use

We first quantified the sex differences in space use and the association between movements and parental behavior across three species that differ in parental sex roles (Figure 1). In *A. femoralis*, where males do most parental care duties, the average male home range was 153 % larger and movement extent 172 % larger than females (Table 1 and S1, Fig. 1). The same pattern was reflected in long-term movements based on capture-recapture data (Annex 1). In *O. sylvatica*, where females perform parental care, the average male home range was 56 % smaller and movement extent 57 % times smaller than females (Fig. 1, Table 1 and S1). The same pattern was reflected in movement extent based on short-term tracking at a different study site (Annex 1). In *D. tinctorius*, where males perform parental care, we found no significant sex differences in home range size or movement extent (Fig. 1, Table 1 and S1) based on tracking data, but males showed wider-ranging long-term movements based on capture-recapture data (Annex 1, Fig. S2). Sex influenced the average daily travel of *A. femoralis*, where the best-fit model included sex, behavior, daytime temperature, and random factors, but not in *D. tinctorius* and *O. sylvatica. Allobates femoralis* males moved more on days of parental care than on mating days (lsm contrast: β = 1.8, P < 0.001) and other days (lsm contrast: β = 1.9, P < 0.001), but equally between mating days and other days (lsm contrast P = 0.75; Fig. 1). *Allobates femoralis* females moved more on days of mating than other days (lsm contrast: β = 0.6, P < 0.001) and were only observed transporting tadpoles once (Fig. 1). *Dendrobates tinctorius* males also moved more on days of parental care than on mating days (lsm contrast: β = 1.3, P < 0.001) and other days (lsm contrast: β = 1.5, P < 0.001), but moved equally between mating days and other days (lsm contrast P = 0.85, Fig. 1). Daily travel of *D. tinctorius* females did not differ on days of mating from other days (lsm contrast P = 0.4, Fig. 1), and females were never observed in the pools or with tadpoles. Daily travel of *O. sylvatica* males did not differ between mating days and other days (lsm contrast P = 0.95, Fig. 1) and males were never seen transporting tadpoles, but were regularly found near the breeding pools often located inside their territories. Females of *O. sylvatica* moved more on days of parental care than on other days (lsm contrast: β = 0.4, P = 0.004; Fig. 1), but there was no difference between mating days and parental days (lsm contrast P = 0.8) or other days (lsm contrast P = 0.3). In summary, we found sex differences in space use that reflect sex differences in parental roles across species, and that parental care is associated with the longest movements in all three species.

**Figure 1.**
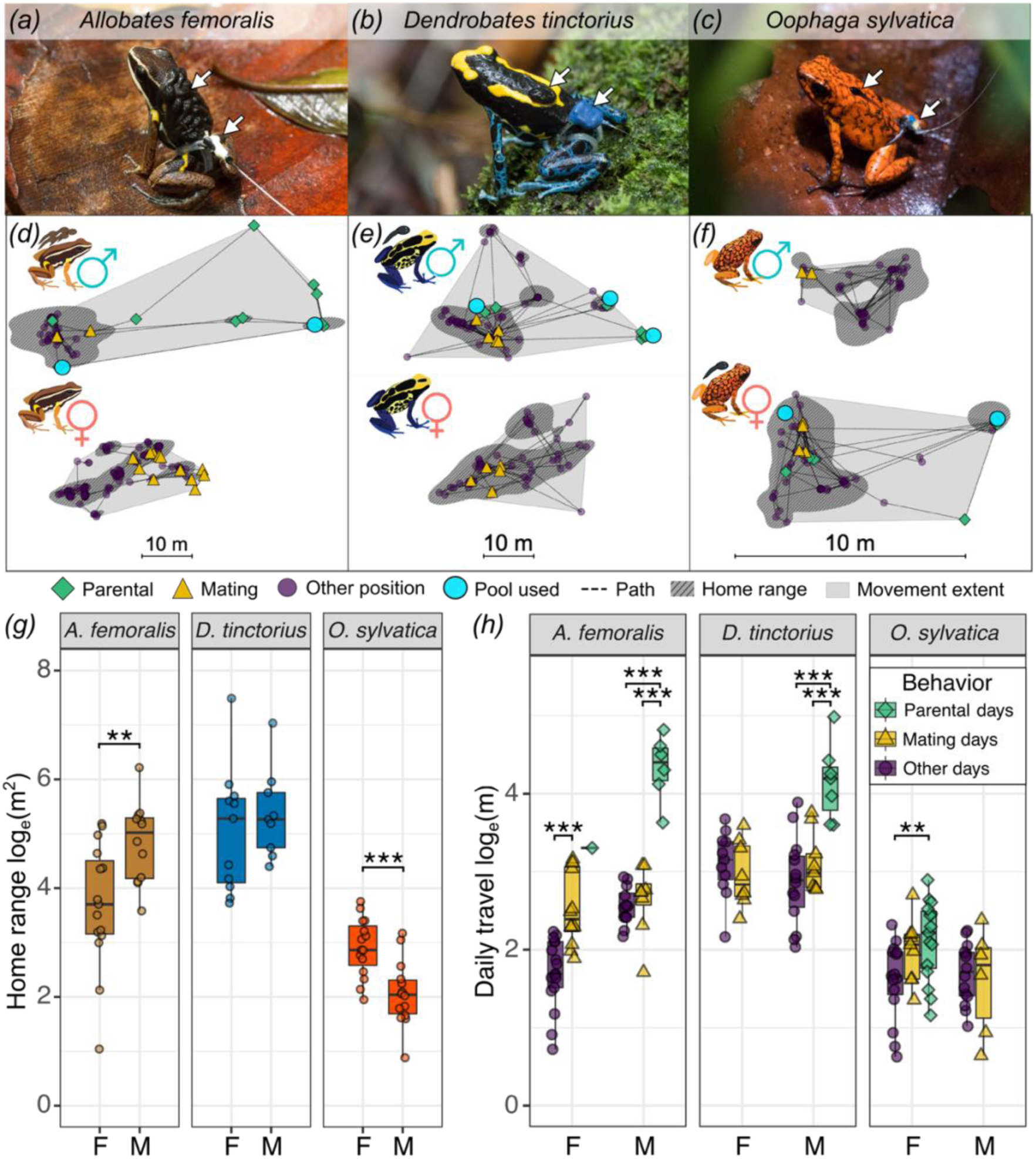
Parental sex roles and behavior drive sex differences in poison frog space use. Male (a, b) and female (c) individuals of each study species transporting tadpoles while wearing a tracking tag. White arrows indicate tadpoles and the tag. (d – e) Examples of representative space use patterns of one individual of each species and sex show different measured space use parameters. We calculated the daily travel as the cumulative distance (line) between all relocations (points) per day; the movement extent (gray shaded area); and the home range representing more intensely used areas (darker hatched area). Frog positions are classified to represent three types of behaviors associated with daily movements: parental behavior (green diamonds), mating behavior (yellow triangles), and other (purple circles). Light blue circles represent pools used for tadpole deposition. Note that the scale is different in the panel for (f) *O. sylvatica*. Boxplots show sex differences in home range size (g), and daily travel (h) between days when parental behavior, mating, or neither were observed. Plot rectangles indicate the lower and upper quartiles with the median line, whiskers extend to 1.5 times the interquartile limited by value range, and dots indicate individuals. As frogs were tracked for multiple days, average values per individual per behavioral category are shown. Days were categorized as pool visits or mating days if the corresponding behavior was observed at least on one relocation of that day. Y-axes are log_e_-transformed. Statistical significance levels are indicated as *p 0.05 – 0.01, **p <0.01, ***p <0.001.

**Table 1.** Home range size model summaries. Summary of three linear models with log_e_-transformed home range size in *A. femoralis*, *D. tinctorius*, and *O. sylvatica* as the response variable, sex as the predictor, and tracking duration (in days) as a covariate. Statistical significance with p < 0.05 is highlighted in bold.

|  | *A. femoralis*<br>log<sub>e</sub> (home range) |  | *D. tinctorius*<br>log<sub>e</sub> (home range) |  | *O. sylvatica*<br>log<sub>e</sub> (home range) |  |
| --- | --- | --- | --- | --- | --- | --- |
| Predictors | Estimates (CI) | *p* | Estimates (CI) | *p* | Estimates (CI) | *p* |
| (Intercept) | 3.15<br>(1.1 – 5.2) | **0.004** | 5.3<br>(4.0 – 6.6) | **<0.001** | 2.5<br>(1.6 – 3.4) | **<0.001** |
| Sex [male] | 0.9<br>(0.2 – 1.7) | **0.015** | 0.2<br>(-0.9 – 1.3) | 0.68 | -0.8<br>(-1.3 – -0.4) | **0.001** |
| Tracking duration | 0.06<br>(-0.1 – 0.2) | 0.46 | -0.02<br>(-0.1 – 0.06) | 0.62 | 0.04<br>(-0.04 – 0.1) | 0.36 |
| Observations | 25 |  | 19 |  | 29 |  |
| R<sup>2</sup> / R<sup>2</sup> adjusted | 0.28 / 0.21 |  | 0.02 / 0.0 |  | 0.40 / 0.35 |  |

### Sex differences in parental care do not predict navigational performance

We tested if there were sex differences in navigational performance that reflected sex differences in parental roles and space use across species. When translocated 50 meters, *A. femoralis* males were more likely to return home (81% males, 44 % females), but both sexes of *D. tinctorius* (94% males, 94% females) and *O. sylvatica* (80% males vs 70% females,) were equally likely to return (Fig. 2 and Table 3). When translocated 200 meters, only males of *A. femoralis* (75% males, 0% females) and both females and males *D. tinctorius* (56% males, 39% females), but none of *O. sylvatica* were able to return home (Fig. 2 and Table 2). One *O. sylvatica* male could not be located after 6 days and was found back home two days later. *Allobates femoralis* were less likely to home back with higher daytime temperature, but the daytime temperature had no effect on homing success in *D. tinctorius* and *O. sylvatica* (Table 2). Frog weight had no effect on homing success. When translocated 50 meters, *A. femoralis* males explored larger areas than females, but we observed no sex difference in *D. tinctorius* and *O. sylvatica* (Fig. 3, Table S2). When translocated 200 meters, males of all three species were more explorative than females (Fig. 3, Table 3). In addition, *A. femoralis* explored less with higher daytime temperature, but the daytime temperature did not affect the explored area in *D. tinctorius* or *O. sylvatica* (Table S2 and 3). Frog weight positively influenced the explored area only in *D. tinctorius* translocated 200 meters (Table 3). Among frogs that returned home, *A. femoralis* males returned more directly and faster from 50 meters when compared to females (Fig. 3, Table 4 and S3). In *D. tinctorius*, females returned in more direct paths than males from 200 m (Fig. 3, Table 4). We found no sex difference in *O. sylvatica* homing. The same sex-difference pattern in homing accuracy was observed based on angular deviation data (Annex 2). *Allobates femoralis* and *O. sylvatica* returned in less direct paths and slower with higher daytime temperature. *Dendrobates tinctorius* returned slower with higher daytime temperature, but temperature had no effect on trajectory straightness (Table 4 and S3). Frog weight did not affect the homing duration.

**Figure 2.**
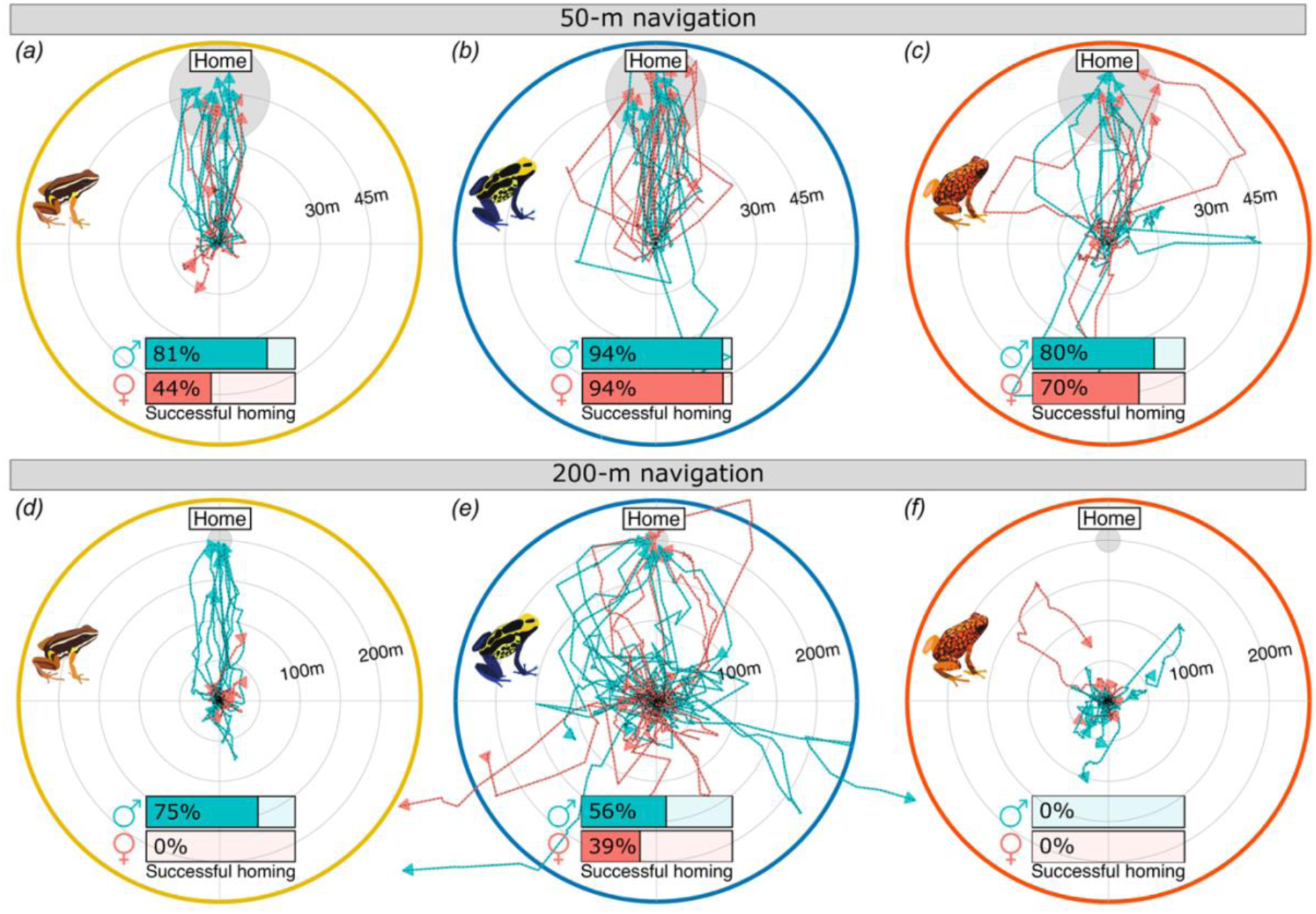
Species and sex differences in movement trajectories of translocated poison frogs. (a - f) Homeward normalized movement trajectories of (a , d) *A. femoralis*, (b, e) *D. tinctorius*, and (c, f) *O. sylvatica* translocated approximately 50 meters (a – c) or 200 meters (d – f) from home. All trajectories are normalized to a common start location (center of the plot) and home direction (top of the plot). The approximate home area is indicated by a gray circle. Each line corresponds to a different individual with male trajectories in teal and female in red. The proportion of each sex that showed homing behavior is indicated on inserted bar plots. Frogs were considered homing if they completed at least 70% of the distance from the release site to the home center within three or six days for 50 m and 200 m, respectively.

**Figure 3.**
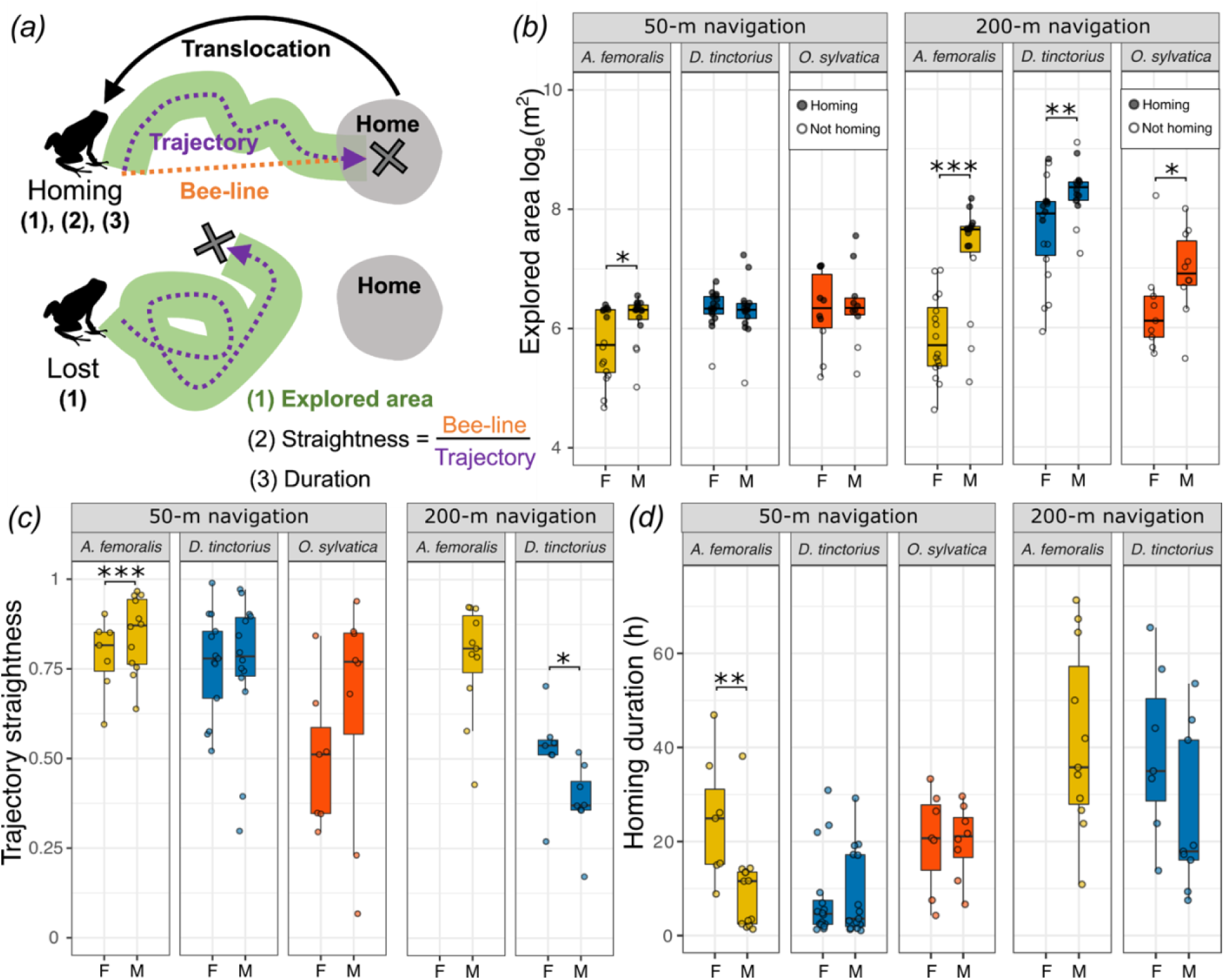
Sex differences in poison frog exploration and navigational performance. (a) Schematic representation of the parameters measured during navigation experiments which are plotted in panels (b - d). Explored area (b), trajectory straightness (c), and homing duration (d) were measured for successful homing, while only explored area (b) were measured for frogs that did not return home. Boxplots show sex differences in (b) explored area (log_e_- transformed), (b) homing trajectory straightness, and (c) homing duration. (a) Filled and empty circles indicate individuals that were homing or not. Plot rectangles indicate the lower and upper quartiles with the median line, whiskers extend to 1.5 times the interquartile limited by the value range, and dots indicate individuals. Statistical significance levels are indicated as *p 0.05 – 0.01, **p <0.01, ***p <0.001.

**Table 2.**
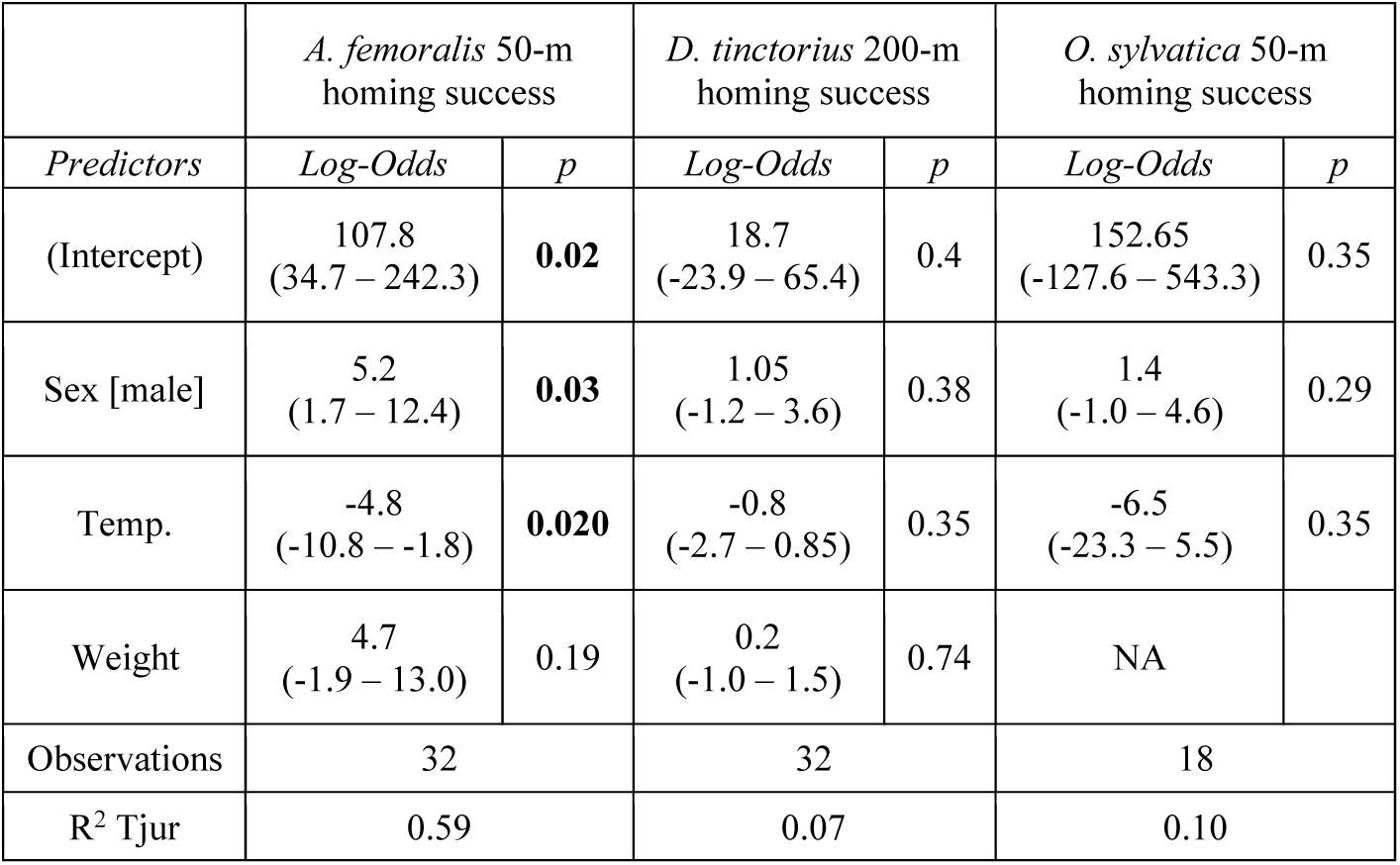
Homing success model summaries. Summary of three logistic regression models with homing success in *A. femoralis*, *D. tinctorius*, and *O. sylvatica* as the response variable, sex as the predictor, and average daytime temperature (Temp.) and frog weight as covariates. We did not perform statistical comparisons conditions *A. femoralis* 200-m because only males successfully returned; for *O. sylvatica* 200-m because no frogs returned and for *D. tinctorius* 50-m because both sexes return at an equal rate. Weight was excluded for *O. sylvatica* to achieve model convergence. Statistical significance with p < 0.05 is highlighted in bold.

**Table 3.** Explored area model summaries. Summary of three linear models with log_e_-transformed explored area in *A. femoralis*, *D. tinctorius*, and *O. sylvatica* as the response variable, sex as the predictor, and average daytime temperature (Temp.) and frog weight as covariates. Statistical significance with p < 0.05 is highlighted in bold.

|  | *A. femoralis* 200-m<br>log<sub>e</sub> (explored area) |  | *D. tinctorius* 200-m<br>log<sub>e</sub> (explored area) |  | *O. sylvatica* 200-m<br>log<sub>e</sub> (explored area) |  |
| --- | --- | --- | --- | --- | --- | --- |
| *Predictors* | *Estimates* | *p* | *Estimates* | *p* | *Estimates* | *p* |
| (Intercept) | 28.0<br>(8.7 – 47.4) | **0.006** | 5.6<br>(-7.65 – 18.9) | 0.39 | 30.85<br>(-97.0 – 158.7) | 0.61 |
| Sex [male] | 1.5<br>(0.9 – 2.1) | **<0.001** | 1.5<br>(0.8 – 2.2) | **<0.001** | 0.8<br>(0.02 – 1.7) | **0.045** |
| Temp. | -1.0<br>(-1.8 – -0.2) | **0.017** | -0.04<br>(-0.6 – 0.5) | 0.87 | -0.9<br>(-6.4 – 4.5) | 0.72 |
| Weight | 1.0<br>(-0.5 – 2.5) | 0.19 | 0.6<br>(0.2 – 0.9) | **0.004** | -1.4<br>(-3.2 – 0.3) | 0.10 |
| Observations | 30 |  | 32 |  | 15 |  |
| R<sup>2</sup> / R<sup>2</sup> adjusted | 0.56 / 0.51 |  | 0.41 / 0.35 |  | 0.38 / 0.21 |  |

**Table 4.**
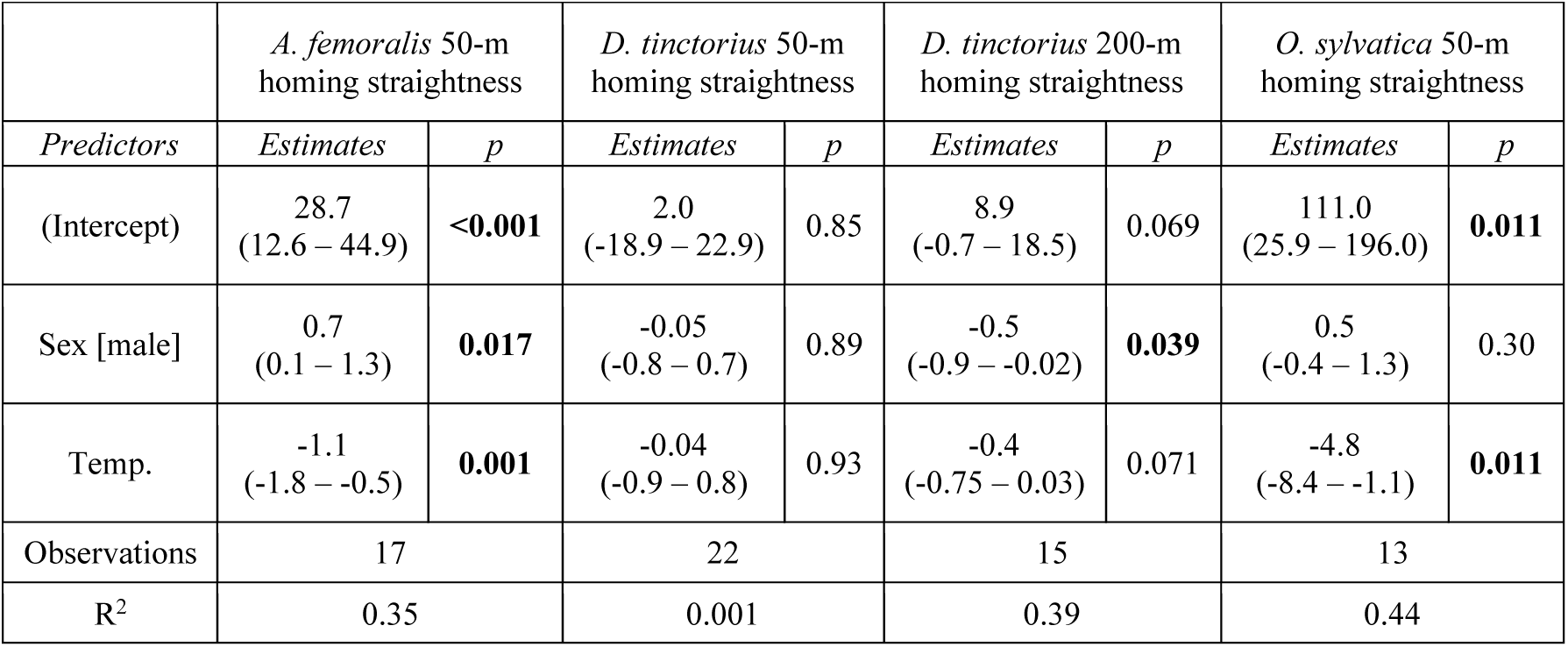
Homing trajectory straightness model summaries. Summary of four beta regression models with homing trajectory straightness in *A. femoralis*, *D. tinctorius*, and *O. sylvatica* as the response variable, sex as the predictor, and average daytime temperature (Temp.) as a covariates. Statistical significance with p < 0.05 is highlighted in bold.

To sum up our navigation experiments, we found that *A. femoralis* males navigate home faster and more accurately, *D. tinctorius* females navigate more accurately from long distances, and males across species display more exploration-related movement compared to females regardless of sex differences in parental care roles.

### Androgens correlate with navigation-associated behavior

We also investigated the relationship between androgen levels and spatial behavior during the navigation task described above. There was a high inter-individual variation of androgen levels in both sexes of all species, but on average, males showed higher androgen levels in all three species (Fig. 4, Table S4). There were no significant differences between baseline and back-home samples in all three species (Fig. 4, Table S4). Baseline levels did not influence exploration, homing duration, nor trajectory straightness in *A. femoralis* (Fig 4, Table S5). Baseline androgen levels, together with sex, translocation distance, and frog weight significantly predicted exploration in *D. tinctorius*, but did not influence homing duration or trajectory straightness (Fig. 4, Table S6). Baseline levels significantly predicted trajectory straightness, but not exploration or homing duration in *O. sylvatica* (Fig. 4, Table S7). The explored area had a significant positive effect and successful homing a significant negative effect on delta androgen levels in *A. femoralis*, but not in *D. tinctorius* and *O. sylvatica* (Table S8).

**Figure 4.**
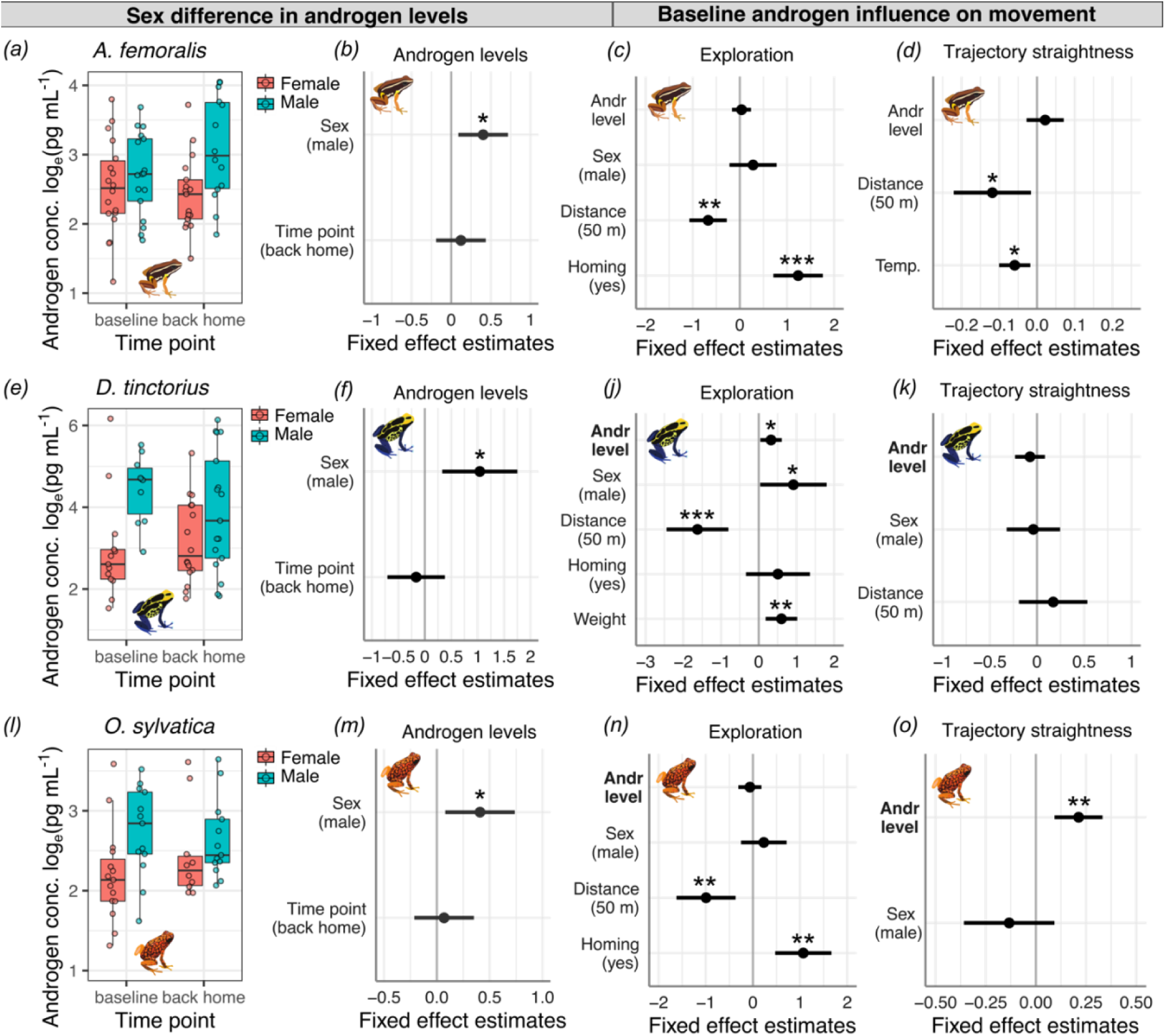
Relationships between androgen levels and spatial behavior. Boxplots show sex differences in water-borne androgen concentration measured before and after the navigational task in (a) *A. femoralis*, (e) *D. tinctorius*, and (l) *O. sylvatica*. The coefficient plots indicate the effect size and confidence intervals of androgen level difference between sexes and the two sampling points for (b) *A. femoralis*, (f) *D. tinctorius*, and (m) *O. sylvatica* and the influence of androgen levels and other factors on exploration (c, j, n) and homing trajectory straightness (d, k, o) in each species. Androgen concentrations are natural loge-transformed. Statistically significant levels are indicated as *p 0.05 – 0.01, **p <0.01, ***p <0.001.

## Discussion

Sex differences in spatial behaviors are typically interpreted through the lens of the adaptive specialization hypothesis, where larger home ranges and better navigational abilities in males are seen as adaptive traits (Jones et al., 2003). This has been countered with the androgen spillover hypothesis, which suggests that enhanced spatial abilities in males are a byproduct of higher male androgen levels rather than an adaptation (Clint et al., 2012). However, there are no comparative studies where females and males of closely related species have reversed spatial behavior and are expected to show a reversal in spatial abilities. Here, we linked the reproductive strategies, space use, navigational performance, and androgen levels in three species of frogs that differ in which sex performs spatially-relevant parental care tasks that tie spatial accuracy to reproductive fitness (Figure 5). We found that parental care shapes sex differences in space use, but no evidence that sex differences in navigational performance are linked to the reproductive strategy. Importantly, we found that females did not outperform males in *O. sylvatica*, the species with more complex female spatial behavior and larger home ranges associated with female parental care. We also found that males of all three species tended to be more explorative than females, and had higher androgen levels. Moreover, increased androgen levels were associated with higher exploration in two species with male care and navigational accuracy in the species with female care, leaving open the possibility that sex differences in spatial behavior might result from sex differences in androgen levels independent of the differences in parental sex roles.

**Figure 5.**
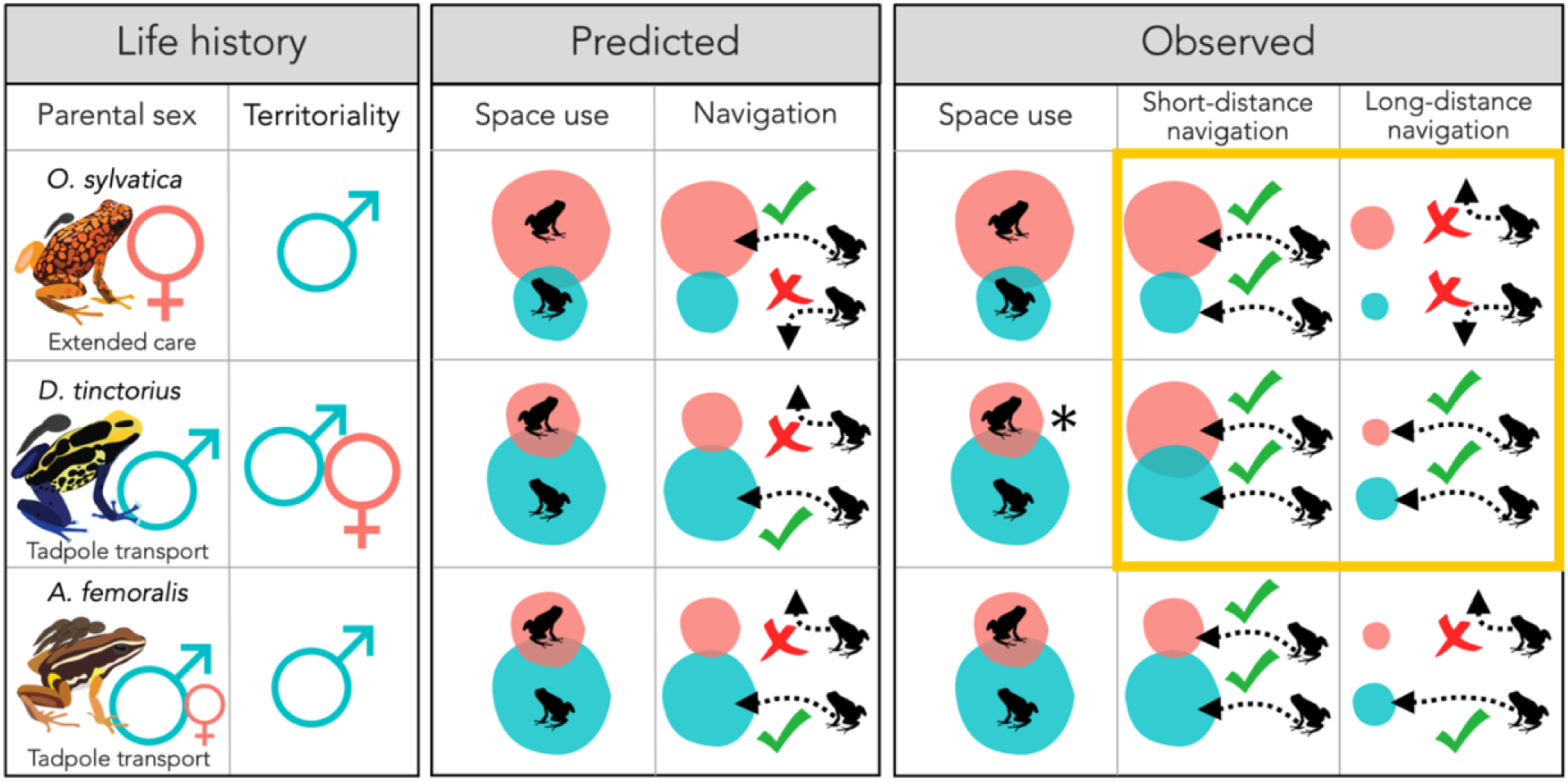
Summary of sex and species differences in life-history traits, predicted, and observed space use and navigational performance. Red and teal circles represent the space use extent for females and males, respectively, and dotted arrows represent homing after translocations. Orange frame highlights the observed outcomes that differed from the predictions. In all cases, we found no sex differences where differences were expected. *Sex difference only observed in long-term movements and qualitatively in vertical space use (see Annex 1).

### Reproductive strategy shapes species and sex differences in space use

We show that parental care in poison frogs increases the mobility of the caring sex and thus shapes the sex differences in space use. In *D. tinctorius* and *A. femoralis*, males transport tadpoles outside their territory and provide no further care. As we predicted, *A. femoralis* males have a much wider space use than females. Male long-distance movements were primarily observed during tadpole transport and occasional territory shifts, while female space use extent was mainly driven by mate-seeking. This sex difference was less pronounced in *D. tinctorius*, where males did not differ from females in short-term space use based on tracking. Males, however, moved over a wider range based on long-term capture-recapture data (Annex 1). The apparent difference between short-term and long-term data in *D. tinctorius* could be due to real sex differences only emerging in longer-term space use patterns, or the limited statistical power to detect a slight sex difference with a much smaller sample size in the tracking dataset. *Dendrobates tinctorius* males also moved the longest distances on tadpole transport days. During tadpole transport, males also climbed vertically to reach water-filled treeholes high above the ground, which they are known to use for tadpole deposition (Fouilloux et al., 2021). Our space use data does not capture these vertical movements and thus might considerably underestimate *D. tinctorius* male mobility and associated sex differences in vertical space use qualitatively and quantitatively. We could not identify what drove wide-ranging horizontal movements in *D. tinctorius* females, but seeking better foraging sites has been suggested among the movement drivers in *D. tinctorius* (Born et al., 2010).

Including a species with female uniparental care whose spatial movements are tied to reproductive fitness allowed us to ask whether enhanced mobility could be an adaptive trait linked to parental care. *Oophaga sylvatica* females must remember and revisit tadpole nurseries dispersed over tens of meters to provision their young with trophic, unfertilized eggs (Stynoski, 2009; Summers, 1992; this study). As predicted by the adaptive specialization hypothesis, tracking revealed larger female home ranges in *O. sylvatica*. Females typically moved between several reproductive pools and moved most when visiting or switching between the pool sites. Like *D. tinctorius* males, *O*.

*sylvatica* females regularly climbed vertically to water-filled plants up to ten meters above the ground, but these vertical movements are not captured in our space use data. Therefore, our data underestimated the mobility of *O. sylvatica* females and the resulting magnitude of the sex difference. In contrast, males did not climb above two meters, and their movements were restricted to exceedingly small calling territories. It remains unknown if males explore larger areas over the long term, particularly when searching for new territories. Overall, *O. sylvatica* showed more restricted movements than the two species with predominantly male parental care. Several previous studies in poison frogs indicate that species with extended female care, which includes tadpole provisioning, show more restricted space use than closely related species with male uniparental care (Brown et al., 2009; Donnelly, 1989; McVey et al., 1981; Murasaki, 2010; Pašukonis et al., 2019; Summers, 1992). Our data confirm that sex and species differences in space use across poison frogs can be explained, in big part, by species differences in the care-providing sex and the intensity of parental care.

Research on sex differences in space use has typically focused on differences in mating strategies. For example, male bias in larger home ranges is well documented in rodents, where polygamous species tend to have greater sex differences than monogamous species (reviewed in Clint et al., 2012; Jones et al., 2003). In lizards, a meta-analysis of home range size in 60 species found that males have larger home ranges than females, and suggested that this sex difference is related to the mating system and access to females (Perry & Garland Jr, 2002). Our study shows that parental care can directly influence the space use of the caring sex resulting in sex differences in space use. Parental care has been hypothesized to reduce mobility in the caring sex because the research has focused on maternal care in lactating mammals (Barnett & McEwan, 1973; Sherry & Hampson, 1997; Trivers, 1972). In contrast, moving with the offspring or for offspring provisioning is taxonomically widespread in vertebrates and invertebrates (Choe et al., 1997; Clutton-Brock, 1991; Kölliker et al., 2012) and can increase mobility in the caring sex, thereby shaping sex differences in space use patterns in taxonomically diverse groups.

### Species differ in navigational and movement strategy

All three species showed well-developed homing ability, which is consistent with previous studies in amphibians, including various anurans (Arcila-Pérez et al., 2020; Dole, 1968; McVey et al., 1981; Navarro-Salcedo et al., 2021, 2022; Pašukonis et al., 2014, 2018; Pichler et al., 2017; Shaykevich et al., 2021; Sinsch, 1987, 1992) and caudates (Diego-Rasilla et al., 2005; D. Grant et al., 1968; Joly & Miaud, 1989; Phillips et al., 1995; Sinsch, 2007; Twitty et al., 1964; reviewed in Ferguson, 1971; Sinsch, 2006; Wells, 2010). Despite limited movement capacity and sedentary lifestyle, many amphibians appear to share a general ability to navigate home after translocations from distances exceeding their routine movements (Sinsch, 1990, 2014). Moreover, the fact that species showing such tremendous variation in life history share this capacity suggests that well-developed navigational abilities play a fundamental role in amphibian reproduction and survival. Like many other tropical anurans, poison frogs rely on moving between small and scattered water bodies for reproduction, which might have selected for particularly highly developed navigational abilities in this group.

Our results indicate that navigational and movement strategies differ even between closely related species. The scale and strategy of navigation varied between species and were related to species differences in home range size and reproductive strategy. For example, *O. sylvatica* did not return from longer translocation distances, in line with their much smaller home ranges than in *A. femoralis* and *D. tinctorius*. The correlation between home range size and homing performance further supports the hypothesis that poison frogs rely on exploration and spatial learning for navigation (Pašukonis et al., 2014, 2016, 2018). We also found notable species differences in movement and search strategies when animals were navigating home after translocations (Fig. 2, Annex 3, Fig. S4). *Allobates femoralis* tended to stay close to the release site for prolonged periods and then navigate back home via a direct path. Non-homing individuals moved very little and remained close to the release site. However, *D. tinctorius* showed wide-ranging exploratory movements and usually returned home through an indirect and lengthy route. Similarly, even though *O. sylvatica* returned home only from shorter distances, they also showed some wide-ranging exploratory behavior that was never observed in *A. femoralis*. We hypothesize that some of these species differences could be linked to movement strategy differences selected under different predation pressure. *Dendrobates tinctorius* and *O. sylvatica* are brightly colored species that forage actively to acquire their alkaloid-based chemical defenses from the invertebrate diet (Santos et al., 2016; Santos & Cannatella, 2011). Aposematic coloration may reduce predation pressure and the cost of movement while potentially increasing exploration (Carvajal-Castro et al., 2021; Pough & Taigen, 1990; Speed et al., 2010; Summers, 2019; Toft, 1981), allowing different navigational strategies. *Allobates femoralis*, on the other hand, is cryptically colored, non-toxic, and a generalist sit-and-wait forager. Predation pressure is likely to be much higher and the movement more costly for cryptic species, thus potentially selecting more efficient orientation strategies. These differences between closely related species provide a remarkable system for future work on the selective pressures shaping the animal movement and navigational strategy.

### Navigational performance provides no evidence for adaptive sex differences

The adaptive specialization hypothesis is the leading hypothesis to explain variation in home range size and spatial memory between species and sexes in mammals. It predicts that adaptive sex differences in navigational ability are linked to life history traits. We found no evidence for adaptive sex differences in navigational ability in poison frogs. Contrary to our prediction, we found no sex difference in the navigational performance of *D. tinctorius*, a species with male uniparental care. Not only did males not outperform females, but females even showed slightly more accurate homing trajectories than males when navigating long distances. However, the lack of sex differences in the navigational performance of *D. tinctorius* somewhat fits the limited sex differences in space use observed in this species. Crucially, although females of *O. sylvatica* have larger space use and perform tadpole transport and egg provisioning, we found no sex differences in the navigational performance. Previous research on amphibians has also shown patterns inconsistent with sex differences in spatial abilities being an adaptive trait linked to reproductive strategy. Place discrimination tasks have not revealed consistent sex differences in *Engystomops pustulusos*, a frog species where females seek for and move between the males (Liu & Burmeister, 2017; Ventura et al., 2019). Using translocation and recapture methods, one recent study in *Andinobates bombetes,* a poison frog with male uniparental care, found no sex differences in homing rates after translocation (Arcila-Pérez et al., 2020). Another recent recapture study in the glass frog *Ikakogi tayrona*, a species with prolonged maternal care and male territoriality, found that only males, but not females, showed homing after translocations (Navarro-Salcedo et al., 2022). Together, our findings and the growing literature on amphibian navigation parallel findings in mammals, where males typically outperform females in spatial tasks or no sex differences are found.

The only species with marked sex differences in navigational performance was *A. femoralis*, where males were more likely to return home, returned from longer distances, and returned faster and more accurately than females. This finding is partially unexpected because *A. femoralis* females commute between males for reproduction (M.-T. Fischer et al., 2020; E. Ringler et al., 2012) and remember the exact locations of their clutches (E. Ringler et al., 2016). However, because non-homing *A. femoralis* females typically stay stationary, it is difficult to disentangle the lack of motivation from their inability to return home. We expected that *A. femoralis* females would be motivated to return home because they show site fidelity (M.-T. Fischer et al., 2020; M. Ringler et al., 2009; this study) and monitor the presence of their mating partners to eventually take over tadpole transport in case of male absence (E. Ringler, Pašukonis et al., 2015). Indeed, females returned, albeit slower from shorter translocation distances, indicating that they were motivated to return in a sufficiently familiar area. In addition, homing females showed less directed homing, suggesting a sex difference in orientation accuracy. However, males may be more motivated to return home quickly because they risk losing both their territory and all current offspring due to territorial takeovers and potential cannibalism by other males (E. Ringler et al., 2017). We believe that male *A. femoralis* likely have better navigational abilities than females, but the motivational state linked to each individual’s current reproductive or parental status may explain some of the sex and inter-individual differences observed in homing performance.

### Males explore more than females

Males are bolder and more explorative in several species and taxa (e.g., fish: Harris et al., 2010; King et al., 2013; bird: Schuett & Dall, 2009) and multiple adaptive hypotheses have been proposed to explain increased exploration (reviewed in Schuett et al., 2010; Trivers, 1972). Sex differences in exploration tendency could be connected to the sex-biased dispersal observed in different mating strategies, where male-biased dispersal is common in polygamous mammals while female-biased dispersal is common in monogamous birds (Greenwood, 1980; Li & Kokko, 2019; Mabry et al., 2013). However, in birds and mammals, sex, mating systems, and parental care are tightly linked, making it difficult to disentangle factors shaping sex differences in exploratory behavior. In the present study, males of all three species tended to be more explorative than females, particularly when translocated longer distances and, therefore, in less familiar environments. Even in *O. sylvatica*, a species where females perform parental care and have wider space use, males tended to be more explorative after translocations. Male-biased dispersal and higher male exploration rates have also been observed in some frogs without parental care (*Engystomops pustulosus*: Lampert et al., 2003; *Bufo bufo*: Ogurtsov et al., 2018; *Xenopus tropicalis*: Videlier et al., 2015). Thus, regardless of parental care strategies, different life histories, and sex differences in home range size, male amphibians tend to be more exploratory, suggesting that other factors, such as male-biased dispersal and high intra-sexual male competition may be associated with the sex difference in exploration.

### Linking androgens to exploration and navigation

Androgens have been linked to spatial abilities in mammals for several decades (Dawson et al., 1975; Galea et al., 1995; Isgor & Sengelaub, 1998; Joseph et al., 1978; Schulz & Korz, 2010; Sherry & Hampson, 1997; Stewart et al., 1975; Williams et al., 1990). Clint et al. (2012) proposed that the often observed male superiority in spatial navigation might be a side effect of sex difference in androgen levels rather than an adaptation to direct selective pressures on males’ spatial abilities. Our results are somewhat in line with this hypothesis as females did not outperform males when expected based on adaptive predictions. Additionally, although we did not observe a correlation between homing success and androgen levels, we found three associations between androgen levels and spatial behavior.

Higher baseline androgen levels predicted more exploration after translocation in *D. tinctorius*, and the amount of exploration during the navigation task was associated with an increase in androgen levels in *A. femoralis*. We also found that males, on average, had higher androgen levels and higher exploration rates despite the variation in the parental sex roles and high inter-individual variation. Exploration underlies the development of most spatio-cognitive abilities (McNaughton et al., 2006; O’keefe & Nadel, 1978), including spatial memory, presumably used by poison frogs for navigation (Beck et al., 2017; Liu et al., 2016, 2019; Pašukonis et al., 2014, 2016, 2019). Therefore, the association of explorative behavior with androgen levels, especially during the development of spatial memory, might have cascading effects on sex differences in spatial behavior and abilities. Quantifying or manipulating androgen levels during ontogeny and learning, rather than during the spatial task performance, might provide a better understanding of the link between individual differences in the navigational performance observed in our study and androgen levels. We also found that baseline androgens correlated with homing accuracy in *O. sylvatica* in both sexes, further supporting a potential link between androgens and navigational performance. While experimental androgen manipulations are needed to understand the interplay between hormone levels and spatial behavior, our findings lend more support to the androgen spillover hypothesis of sex differences in spatial cognition.

## Conclusions

We found that parental behavior drives space use patterns but not the navigational performance in poison frogs. Most observed sex differences indicated more developed navigational ability and increased exploratory tendency in males, even in species where females show wider-ranging movement for parental care and mate-seeking. Indeed, most previous literature on sex differences in vertebrate spatial abilities shows either no sex differences or better performance in males. We also found higher average androgen levels in males of all three species despite the marked species differences in parental sex roles and aggressive female behavior. However, there was a high inter-individual variation in androgens and a large overlap between sexes. Some of this inter-individual variation in androgen levels was related to individual differences in exploration and homing accuracy, suggesting an interplay between androgens and spatial behavior. Therefore, our findings are more consistent with the androgen spillover hypothesis than the widely accepted adaptive specialization hypothesis. We speculate that sex differences in spatial abilities could be a byproduct of selective pressures on sexual traits such as aggressiveness and the associated increase in androgen levels. However, the indirect effects of androgens, such as increased male exploration tendency, are likely to be adaptive and maintained in the context of male territoriality and widespread male parental care in poison frogs.

## Materials and Methods

### Study species

We studied three poison frog species with different life histories and parental sex roles: the Brilliant-Thighed Poison Frog (*Allobates femoralis* [Boulenger 1884], Aromobatidae), the Dyeing Poison Frog (*Dendrobates tinctorius* [Cuvier 1797], Dendrobatidae), and the Diablito Poison Frog (*Oophaga sylvatica* [Funkhouser 1956], Dendrobatidae). *Allobates femorali*s and *D. tinctorius* occur in syntopy in the Guiana Shield, but *A. femoralis* has a wider range across Amazonia (T. Grant et al., 2006). *Oophaga sylvatica* is endemic to the Chocoan Rainforest of the Pacific Coast of Ecuador and Colombia (T. Grant et al., 2006). All three species are diurnal, breed throughout the local rainy season, and shuttle tadpoles from terrestrial clutches in the leaf litter to aquatic tadpole nurseries (T. Grant et al., 2006; Silverstone, 1975, 1976). In *A. femoralis*, groups of up to ∼ 20 tadpoles are predominantly transported by territorial males and deposited in terrestrial pools (E. Ringler et al., 2013; Roithmair, 1992). Females take over tadpole transport when males disappear (E. Ringler, Pašukonis, et al., 2015). In *D. tinctorius*, one or two tadpoles are transported by males (but see E. K. Fischer & O’Connell, 2020; Rojas & Pašukonis, 2019 for reports of female transport) to terrestrial and arboreal pools (Fouilloux et al., 2021; Rojas, 2014, 2015; Rojas & Pašukonis, 2019). After tadpole deposition into pools, no further parental care is provided in *A. femoralis* and *D. tinctorius*. In contrast, one or two tadpoles of *O. sylvatica* are transported by females and deposited into water-filled plants (Silverstone, 1973; Summers, 1992; Zimmermann & Zimmermann, 1981). Tadpoles feed on unfertilized eggs, which the mother returns to provide every ∼ 3 - 7 days (this study and personal observation by E. Tapia communicated to LAC). In all three species, males and females show site fidelity, but the levels of aggressiveness and territoriality vary. In *A. femoralis* and *O. sylvatica*, males vocally advertise and aggressively defend small territories, while females visit males for mating and show no aggressive behavior (M.-T. Fischer et al., 2020; M. Ringler et al., 2009; Roithmair, 1992; Silverstone, 1973; Summers, 1992; this study). In *D. tinctorius*, both males and females show intra-sexual aggression as part of territoriality and/or mate guarding, but males do not vocally advertise(Born et al., 2010; Rojas & Pašukonis, 2019; this study).

### Study sites

Data for *A. femoralis* and *D. tinctorius* were collected over five field seasons between 2016 and 2020 in two different plots at the Nouragues Ecological Research Station (4°02’ N, 52°41’ W) in the Nature Reserve Les Nouragues, French Guiana. One plot consists of ∼25 ha of lowland rainforest bordering the Arataï river where both *A. femoralis* and *D. tinctorius* naturally occur. The other plots consist of a 5-ha island in the Arataï river where *A. femoralis* and *D. tinctorius* were absent, but an experimental population of *A. femoralis* was successfully introduced in 2012 (E. Ringler, Mangione, et al., 2015). The island population relies primarily on an array of artificial pools for breeding but otherwise lives under natural conditions (E. Ringler et al., 2018). Data for *O. sylvatica* were collected at two sites in Ecuador: Sapoparque La Florida (0°15’ S, 79°02’ W) in 2017 and Reserva Canandé (0°32’ N, 79°13’ W) in 2019. The La Florida study area (enclosure site hereafter) consisted of a free-ranging population of *O. sylvatica* introduced and kept in two forest enclosures of ∼0.25 ha each inside their natural habitat. The natural breeding pools in the enclosures were supplemented by a high density of small artificial plastic pools and suitable plants. The Reserva Canandé study site (natural site hereafter) consisted of a natural *O. sylvatica* population, relying only on natural pools (water-filled plants). For summarized study site and different dataset information see Annex 6 (Tables S9 and S10)

### Frog tracking

To quantify frog movements, we used two tracking methods, which have been previously used to track poison frogs: harmonic direction-finding and radio-tracking (for more details see Annex 5). We tagged and tracked 311 frogs, located each frog multiple times a day (see further below), and recorded its position and any observed behavior. All study plots were mapped using precision compasses and laser distance meters, establishing a network of labeled reference points (for method details, see M. Ringler et al., 2016). The digital maps were used with the GIS software ArcPad 10 (ESRI, Redlands, USA) on handheld devices for recording the data in the field. Occasionally, when frogs moved out of the mapped area, we recorded their location by GPS position averaging. All data were collected as GIS spatial points with associated behavioral information, checked point-by-point at least twice, and corrected for errors such as wrong frog identities, duplicates, and impossible locations by one of the experiments. All suspect points where frog identity or location could not be unambiguously confirmed and corrected were removed.

### Quantification of space use

To quantify space use, we tagged 36 *A. femoralis*, 31 *D. tinctorius*, and 31 *O. sylvatica*, which we localized 6055 times and tracked for periods ranging from <1 to 45 days per individual. A subset of these data was used in previous publications on female space use in *A. femoralis* (M.-T. Fischer et al., 2020) and tadpole transport in *D. tinctorius* (Pašukonis et al., 2019). When a frog lost its tag, we attempted to recapture and retag the individual to continue tracking. Identity was confirmed based on unique dorsal or ventral coloration patterns. We excluded all frogs that were tracked for less than two full days (not including the tagging and tag removal/loss day) without any manipulation and handling (tag checks or retagging, see Annex 5). We also removed short and temporally disconnected tracking periods when the frog was retagged. In the end, we had data to quantify the space use of 29 *A. femoralis* (17 females and 12 males) tracked from 5 to 16 days (median = 14 days) and 26 *D. tinctorius* (11 females and 15 males) tracked for 3 to 45 days (median = 14 days). We tracked 29 *O. sylvatica* (14 females and 15 males) for 7 to 20 days (median = 10 days) within enclosures and an additional 37 *O. sylvatica* (20 females and 17 males) for 2 to 8 days (median = 3 days) at the natural site.

The sampling rate (number of locations per day) varied for different datasets and tracking periods. In particular, movements associated with specific behaviors of interest, such as the tadpole transport, were often sampled at a higher frequency. As the sampling rate influences the spatial parameters, we standardized the data by down-sampling all datasets to match the datasets with the lowest sampling rate (∼ 4 points per day). An experienced observer (AP) down sampled the data in a two-step procedure that allowed to maintain the maximum spatial and behavioral information. We first counted the number of points per day and selected the days with more than 2 points above the daily average for the respective dataset. We then removed redundant points while trying to keep the most spatially and temporally distributed points, and points with rare behaviors, such as parental care and mating. In a second step, we automatically down-sampled the remaining dataset to a minimum sampling interval of 60 minutes while retaining intermediate points for long (>20 m), fast movements, occurring within shorter than 60 minutes. The resulting dataset had 3 to 7 points per day per frog (median = 4).

For summary of space use variables see Annex 6 (Tables S11 and S12). To quantify the space use, we calculated the daily cumulative distance traveled (daily travel hereafter) for 84 frogs tracked for at least two full days. For 76 frogs tracked for a minimum of seven days, we also quantified the maximum movement extent area (movement extent hereafter) as a minimum convex polygon (MCP) and the home range as 95 % utilization density (UD) contour derived from kernel density estimation (KDE) using the “mcp” and “kernelUD” functions of the “adehabitatHR” R package (Calenge, 2006). The smoothing parameter for the KDE was calculated with a conservative plug-in bandwidth selection method with the “hpi” function of the “ks” R package (Chacón & Duong, 2018).

To quantify the influence of parental and reproductive behaviors on movement, we grouped behaviors into three categories: parental and pool associated behavior (parental behavior hereafter), mating associated behavior (mating behavior hereafter), and “other” when neither parental nor mating behavior was observed. Under parental behavior, we included direct observations of tadpole transport and egg-feeding, as well as all points where the frogs were located within one meter of known breeding pools. Males of *O. sylvatica* were sometimes observed next to breeding pools, but we did not consider that as parental behavior as parental care has not been reported in male *O. sylvatica* nor observed in this study. Under mating behavior, we included direct observations of courting and mating behaviors, as well as all points where the frogs were located within one meter from an opposite-sex individual. We categorized each tracking day as “parental”, “mating”, or “other” whenever the respective behavior was observed at least once on that day. On the days when both parental and mating behavior occurred (12 out of 84 parental behavior days), we categorized the day as “parental” because parental movements were larger in scale. To evaluate the influence of behavior on movement, we then compared the distance traveled on “parental”, “mating”, and “other” days.

Our space use measures represent a snapshot of an animal’s long-term movement, and some species were tracked in experimental study plots confined by enclosure or water. Therefore, we further validated our tracking data described above with three supplementary datasets, including short-term tracking of *O. sylvatica* in a natural population and multi-year capture-recapture data in natural populations of *A. femoralis* and *D. tinctorius* (Annex 1).

### Quantification of navigational performance

To quantify the navigational performance, we carried out translocations and measured homing for 64 *A. femoralis* (32 females and 32 males), 67 *D. tinctorius* (35 females and 32 males), and 39 *O. sylvatica* (19 females and 20 males). Most frogs were tagged and tracked for at least 24 hours before translocation to establish site fidelity to the tagging areas. We presumed the tagging area to correspond to defended territories or core areas within the home range (collectively termed home areas hereafter). We further confirmed site fidelity by behavioral observations of calling, courtship, and repeated use of shelters. In a few instances, *A. femoralis* were translocated immediately after tagging because the territories were already known from a concurrent study (Rodríguez et al., 2020). We did not translocate frogs transporting tadpoles or continuously moving away from the tagging site. The locations of each frog recorded in the home area before the translocation were used as reference points to establish the correct homeward direction and homing success. We translocated one male and one female simultaneously 50 or 200 meters away from their respective home areas. Frog movements were then tracked for 4 or 6 days for 50-m and 200-m translocation, respectively, or until the frogs returned within ∼10 meters from their home areas. Frogs that did not return home within the given time were captured and released back at their home areas. In a few instances, the frogs were tracked for longer than 4 or 6 days, but the trajectories were truncated for the analysis.

For translocation, each frog was captured and placed in an airtight and opaque container. We chose, measured, and located a release site on the study area map using a portable GIS/GPS device and carried the frogs to the release site. To disorient the frogs, the container was rotated multiple times and never carried in a straight line from the capture to the release site. All frogs were captured and released in the afternoon, and the translocation usually took between 30 to 90 minutes. We translocated one male and one female simultaneously from the same area towards the same direction and attempted to vary the translocation direction between pairs within the landscape constraints of the field site. Most frogs were released by placing the container on the ground for at least 5 minutes and then gently opening the lid allowing the frog to leave. In the case of *A. femoralis* tracked in 2017, frogs were removed from a bag by hand and placed under a flower pot with an opening for an exit. If the frogs did not leave the pot within 30 to 60 min, we lifted the pot by hand. These periods of 30 to 60 min were not included in the analysis.

We directly observed and mapped the initial movements of each frog for at least 30 min after release. For this, we set up a radial grid with a 3-meter radius at the release site, made of colored strings for visual reference, and mapped the frog movements at approximately 0.3 m precision on a tablet PC running a custom Python (version 2.7.3) script allowing us to record the frog position in relation to the visual reference grid. Following the direct observation, we attempted to locate the frogs approximately every 20 to 60 minutes, although longer sampling gaps occurred due to local terrain and weather constraints.

For summary of navigation variables see Annex 6 (Tables S11 and S12). To measure the movement strategy and the navigational performance, we quantified (1) homing success, (2) explored area, (3) homing trajectory straightness, (4) homing duration, (5) and angular deviation from the home direction (Annex 2). We assumed that the frog is showing homing behavior (yes/no) if the frog approached at least 70% of the distance from the release site to the home area center, defined as the geometric average of frog positions prior to translocation. To estimate the explored area, we calculated the total area within five meters around the movement trajectory. The value is based on a putative perceptual range of a frog being at least five meters. Trajectory straightness was calculated as the ratio between the straight-line distance from the release site to the end of the homing trajectory and the cumulative distance of the actual homing trajectory. To calculate explored area and trajectory straightness, trajectories were down-sampled to the minimum sampling interval of 15 minutes. The homing duration was calculated as the time from the release of the frog to the moment when the frog crossed a 10-meter buffer drawn around the home area polygon. Night-time (12 hours per night) was excluded from homing duration because none of the study species moves at night. Homing success, explored area, and angular deviation were calculated for all translocated frogs. Trajectory straightness and duration were only estimated for the frogs that successfully returned home. Trajectory straightness and angular deviation could not be estimated for some frogs due to missing data in the trajectory.

### Quantification of androgen levels

For a subset of the frogs used in navigation experiments, we quantified androgen levels using non-invasive water sampling, following the methodology described elsewhere (Baugh & Gray-Gaillard, 2021; Rodríguez et al., 2022). Androgen levels were quantified once in the morning within 2 days before translocation (baseline hereafter) and again in the morning after the frogs returned home by themselves or were returned by the experimenter. We did not quantify androgen levels for the *A. femoralis* and *D. tinctorius* translocated in 2017. Each frog was placed in 40 mL of distilled water inside a small glass container with a dark cover for 60 minutes at the frog capture location and released immediately after. The water sample was pushed through a C18 cartridge (SPE, Sep-Pak C18 Plus, 360 mg Sorbent, 55 - 105 μm particle size, #WAT020515, Waters corp., Milford, MA, USA) with a 20 mL sterile syringe. Cartridges were immediately eluted with 4 mL of 98 % EtOH into 5 mL glass vials and were stored at first at 4 C when in the field and then transferred to -20 C until analysis.

Before the ELISA, 1 mL or 2 mL of the original 4 mL sample was dried down with N2 at 37 C and resuspended with 250 uL of the assay buffer (provided in the kit), and incubated overnight at 4 C. Samples were brought to room temperature and shaken at 500 rpm for 1 h prior the assay. Samples were plated in duplicate and assays were performed following the manufacturer’s protocol. Plates were read at 405 nm, with correction at 570 nm, using a microplate reader (Synergy H1, BioTek Instruments, Winooski, VT, USA) and the concentration of androgens was calculated using the software Gen5 (version 3.05, BioTek Instruments, Winooski, VT, USA). The detection limit for the assay was 7.8 and 2000 pg mL-1 and samples that fell out of this range were removed from the analysis. The cross-reactivity of the testosterone antibody with other androgens is below 15% according to the manufacturer’s manual. Samples with the average intra-assay coefficient of variation (CV) above 15% were excluded from the analysis and the resulting average CV was 5.7%. The average inter-assay CV was 7% for five out of nine assays and not available for the other four due to experimenter error. The final reported concentrations were adjusted for the sample volume taken (1 or 2 mL) and sample concentration during drying-resuspending.

### Weather variables

We measured the understory ambient temperature with temperature-data loggers (HOBO U23 Pro v2, Onset Computer Corp, Bourne, USA) placed ∼30 cm above ground and recording temperature at 15 or 30-minute intervals. Because our study species are exclusively diurnal, we measured daytime temperature by averaging all measures from sunrise to sunset. At the *O. sylvatica* enclosure site, we manually measured temperature three times per day at the start, middle, and end of each tracking session with a handheld electronic thermometer (GFTH 95, GHM Messtechnik, Regenstauf, Germany) held slightly above the ground. We averaged the three measurements to obtain daytime temperature. We also calculated mean daytime temperatures for each frog during the navigation experiments by averaging daytime temperatures over the tracking period of each frog (also see Annex 6 Table S12). We did not use the rainfall data because they were strongly correlated with daytime temperature and missing for some tracking periods.

### Data analyses

All statistics were generated in R Studio (version 1.0.153, RStudio Team, 2020) running R (version 3.6.3, R Core Team, 2020). Space-use plots were generated in QGIS (version 2.14, QGIS.org, 2022), box plots, trajectory plots, and bar plots with ggplot2 R package (Wickham, 2016), circular plots with circular R package (Agostinelli & Lund, 2022), model plots with sjPlot R package (Lüdecke, 2021). Schematic representations and further editing were done with Adobe Illustrator (version 25.2.3, Adobe Inc., Mountain View, CA, USA), and Microsoft PowerPoint (version 16.54, Microsoft Corp., Redmont, WA, USA). All variables and statistical models are summarized in Annex 6 (Tables S11, S12, S13)

### Space use

Spatial variables (daily travel, movement extent, home range, and explored area) approximately followed a log-normal distribution and thus we transformed the raw data by natural logarithm to fit the model assumptions. To investigate the sex differences in movement extent and home range, we fitted two linear models (LMs) with movement extent and home range as responses in each model respectively, and sex, species, and their interaction as predictors. As there was a strong interaction between species and sex for both models, we fitted a separate LM for each species with sex as the predictor. As movement extent and home range often correlate with tracking duration, we included the number of days tracked as a covariate. To investigate the influence of behavior and sex on the daily movement, we fitted linear mixed-effects models (LMMs) with daily travel as the response variable using the “lmer” function within the lme4 R package (Bates et al., 2015). We first fitted an LMM for each species with sex, behavior, and daytime temperature as fixed factors and frog identity and tracking date as random factors. We checked for a correlation between daily travel and number of points per day for each species (significant for *D. tinctorius* and *O. sylvatica*, but not for *A. femoralis*) and used number of points per day as a covariate in LMMs when the correlation was significant. For model selection we followed an information-theoretic approach (Burnham & Anderson, 2002) based on the Corrected Akaike’s Information Criterion (AICc). We calculated models with all combinations of fixed factors (behavior, sex, daytime temperature) while keeping random factors (id and date) and the covariate (number of points per day). We selected the best single model using the “model.sel” function within the MuMIn R package (Bartón & Barton, 2020). Because parental behaviors mostly occurred in one sex, we also analyzed the influence of behavior on movement for each sex separately. We fitted separate LMMs for each sex of each species with behavior as a fixed factor, (factor with 2 or 3 levels: parental, mating, and other as appropriate for each species and sex), and frog identity and tracking date as random factors. The daytime temperature was only included as a fixed factor if the best model based on AICc included temperature (it did for *A. femoralis*, but not for *D. tinctorius* and *O. sylvatica*). We used least-squares means contrasts to compare daily travel between behavioral categories. Post hoc comparisons were done with the “emmeans” function within the emmeans R package (Lenth, 2022) with P-values adjusted by Tukey’s method for multiple comparisons (Tukey, 1977).

### Navigational performance

As species and frogs translocated to different distances showed qualitatively very different movement and homing patterns, we fitted separate models per species and per translocation distance. To investigate the sex differences in the homing success we fitted generalized linear models (GLMs) with binomial error distribution and Logit link function using the “glm” function within R package stats (R Core Team 2020). We used the homing success as a binary response variable with sex, frog weight (except for *O. sylvatica*), and the mean daytime temperature during the entire tracking period as predictors. To investigate sex differences in exploration, we fitted an LM with the natural log-transformed explored area as the response variable and sex, mean daytime temperature, and frog weight as predictors. To investigate sex differences in homing trajectory straightness we fitted a beta regression model for proportions via maximum likelihood using the “betareg” function within the R package betareg (Cribari-Neto & Zeileis, 2010). We used straightness (ratio between 0 and 1) as the response variable with sex and mean daytime temperature as predictors. To investigate sex difference in homing duration we fitted an LM with natural log-transformed homing duration as the response variable and sex, mean daytime temperature, and frog weight as predictors.

### Androgen levels and navigation

To investigate the relationship between androgens, sex, and movement, we fitted a series of separate models for each species. We first fitted LMMs with log-transformed androgen levels as the response variable, sex, and time point (two-level factor: baseline or back home) as fixed factors, and frog identity as a random factor. For all three species, there was no interaction between sex and sampling time point, and the interaction factor was excluded. To investigate the relationship between baseline androgens and spatial behavior during navigation, we first fitted an LM for each species with the log-transformed explored area as the response variable and baseline androgen levels, sex, translocation distance, homing success, mean daytime temperature, and frog weight as predictors. For successfully homing frogs, we also fitted two GLMs (Gamma distribution, inverse link function) with trajectory straightness and homing duration as response variables and baseline androgen levels, sex, translocation distance, mean ambient temperature, and frog weight as predictors. To be able to evaluate the relative influence of different fixed factors on the response variables, we standardized all continuous predictors by centering and scaling using the “stdize” function from the MuMIn R package. We did not include sex for GLMs of trajectory straightness and homing duration of *A. femoralis* because androgen levels were available for only one homing female. If movement itself had an influence on androgen levels, we expected the amount of exploration to have a significant influence on androgen levels after the navigation task. Therefore, we calculated delta androgen levels by subtracting the baseline from the back home levels and fitted an LM with delta androgen levels as a response variable and the explored area, sex, tracking duration, homing success, mean daytime temperature, and frog weight as fixed factors. To reduce the number of covariates, for each model mentioned above, we compared the full models against a model without the mean temperature or the frog weight using the “drop1” function of the stats R package. We removed mean temperature and frog weight from the final model if the model excluding these factors (separately) was not significantly different (P > 0.1).

### Ethics statement and permits

We strictly adhered to the current US, French, Ecuador, and European Union law, and followed the ‘Guidelines for use of live amphibians and reptiles in the field and laboratory research’ by the Herpetological Animal Care and Use Committee (HACC) of the American Society of Ichthyologists and Herpetologists (Beaupre et al., 2004); and the Association’s for the Study of Animal Behaviour (ASAB) ‘Guidelines for the use of live animals in teaching and research’ (ASAB, 2020). The research was approved by the Institutional Animal Care and Use Committee of Stanford University (protocol ID 33211, issued to LAO) and by the Animal Ethics and Experimentation Board of the Faculty of Life Sciences, University of Vienna (approvals No 2016-002, 2016-003, 2018-10, issued to AP). The permits in French Guiana were issued by the local authorities (DIREN premits: R03-2016-10-21-002, N°2015-289-0021, n°2011-44/DEAL/SMNBSP’, issued to ER). In addition to these permits, all protocols for fieldwork were approved by the scientific committee of the Nouragues Ecological Research Station (approval communicated to AP) and the Nouragues Nature Reserve (partnership agreement No 01-2019 with AP, BR, ER, MR). The permits in Ecuador were issued by the local authorities (Ministerio de Ambiente, approval document No 013-18-IC-FAU-DNB/MA, issued to LAC). In addition, the authorization to work in Reserva Canandé was given by reserve authority Fundación Jocotoco, Ecuador (approval communicated to AP).

## Data availability

All data and associated scripts will be made available after publication.

## Supporting information

Annex 1

Annex 2

Annex 3

Annex 4

Annex 5

Annex 6

## Acknowledgments

We are grateful to the staff of the CNRS Nouragues Ecological Research Station, Nouragues Nature Reserve, Fundación de Conservación Jocotoco, Reserva Canandé, Centro Jambatu de Investigación y Conservación de Anfibios, and Wikiri for logistic support and generously providing access to the study sites. We are deeply grateful to these organizations and their dedicated staff for their commitment to preserving our natural world and facilitating research. We thank Kristina Beck, Steffen Weinlein, Susanne Stückler, Eva K Fischer, Italo Tapia, and Vincent Premel for assistance in the field, Jinook Oh for coding the FroggerLogger Python script, and Walter Hödl for continuing inspiration and mentorship that has led to this study. This work is part of a partnership agreement between AP, BR, MR, ER and the Nouragues Nature Reserve to improve and spread the knowledge about amphibians.

## Author contributions

AP: Conceptualization (lead), methodology (lead), investigation-field (lead), investigation-laboratory (lead), data curation (lead), formal analysis (lead), data visualization (lead), writing-original draft (lead), funding acquisition (lead), permit acquisition (lead), other resources (lead), supervision (lead).

SJSR: Investigation-field (lead), data curation (supporting), formal analysis (supporting), writing-review & editing (supporting).

MTF: Investigation-field (supporting), data curation (supporting), writing-review & editing (supporting).

MCL: Methodology (supporting), investigation-field (supporting), formal analysis (supporting), funding acquisition (supporting), writing-review & editing (supporting).

DAS: Investigation-field (supporting), data curation (supporting), writing-review & editing (supporting).

BR: Methodology (supporting), investigation-field (supporting), data curation (supporting), funding acquisition (supporting), writing-review & editing (supporting).

MR: Methodology (supporting), investigation-field (supporting), data curation (supporting), writing-review & editing (supporting), other resources (supporting).

ABR: Investigation-field (supporting), writing-review & editing (supporting). AML: Investigation-field (supporting), writing-review & editing (supporting).

ER: Writing-review & editing (supporting), funding acquisition (supporting), permit acquisition (lead), other resources (supporting), supervision (supporting).

CR: Methodology (supporting), investigation-laboratory (supporting), formal analysis (supporting), writing-review & editing (supporting).

LAC: Writing-review & editing (supporting), permit acquisition (lead), other resources (supporting).

LAO: Conceptualization (lead), methodology (supporting), data visualization (supporting), writing-original draft (supporting), writing-review & editing (supporting), funding acquisition (lead), permit acquisition (lead), other resources (lead), supervision (lead).

