## Supplementary material for "Contrasting parental roles shape sex differences in poison frog space use but not navigational performance": Annex 1

### Space use data validation

Our space use data obtained by tracking had several limitations. First, tracking represents only the short-term space use patterns. Second, *A. femoralis* and *O. sylvatica* were tracked in a semi-experimental population supplemented by artificial breeding pools and confined by a natural river barrier (~5 ha island) or artificial enclosure fence (two enclosures of ~0.25 ha each). We, therefore, validated our space use data obtained by tracking data with three supplementary datasets: short-term tracking of *O. sylvatica* at a natural population and multi-year capture-recapture data at a natural population of *A. femoralis* and *D. tinctorius* (for more details on the capture-recapture method see Ringler et al., 2009) for more details on the capture-recapture methods). For *O. sylvatica*, we checked if the sex differences seen in enclosures were consistent with short-term tracking (one to seven days, median = 3 days) of 37 *O. sylvatica* (17 females and 20 males) data collected at a natural population. For both datasets (enclosures and natural site) we measured the movement extent distance, as the maximum linear distance in meters between two locations of the same individual. We then fitted a separate LM for *O. sylvatica* natural site and enclosure data with movement extent distance as the response variable and sex and tracking duration as predictors. We also fitted an LMM for the natural site and enclosure data with daily travel as the response variable, sex as a fixed factor, and frog identity as a random factor. There was no sex difference in the *O. sylvatica* daily travel at the natural site (LMM: $\beta_{\text{daily male}} = -0.27$, $t = -1.6$, $n = 93$, $P = 0.1$) nor at the enclosures (LMM: $\beta_{\text{daily male}} = -0.11$, $t = -1$, $n = 287$, $P = 0.3$; Fig. S1a), but short-term movement extent distances were larger in females than in males of *O. sylvatica* at both the natural site (LM: $\beta_{\text{extent male}} = -0.38$, $t = -2.4$, $n = 37$, $P = 0.02$) and at the enclosure (LM: $\beta_{\text{extent male}} = -0.42$, $t = -2.9$, $n = 29$, $P = 0.007$; Fig. S1b). Therefore, we consider that our tracking data obtained in enclosures is consistent with movement patterns observed in a natural population.

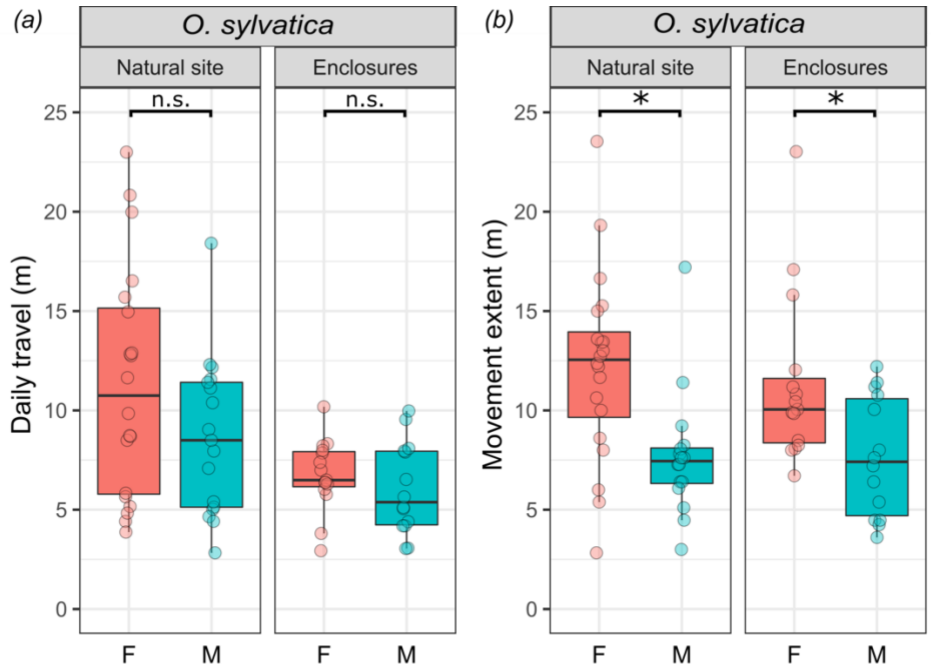


**Supplementary figure S1.** Comparison of sex differences in daily travel and movement extent of *O. sylvatica* tracked in enclosures and a natural population. Boxplots showing sex differences in (a) daily travel and (b) movement extent between female (F) and male (M) frogs tracked at a natural site (left panel) and enclosures (right panel). Plot rectangles indicate the lower and upper quartiles (with the median as the line), whiskers extend to 1.5 times the interquartile range but do not extend past the full value range, and dots indicate individuals. Statistical significance levels are indicated as *p 0.05 – 0.01, **p <0.01, ***p <0.001.

For *A. femoralis* and *D. tinctorius*, we compared the sex differences observed in the tracking data to a multi-year capture-recapture data collected at the same study site over three (2009 – 2011 for *D. tinctorius*, B. Rojas, M. Ringler unpublished data) or six years (2014 – 2019 for *A. femoralis*, M. Ringler, E. Ringler, A. Pašukonis unpublished data). We only included recapture points that were at least 30 days apart for this analysis. For each recaptured individual, we calculated the maximum linear distance between recapture points and compared these distances with the maximum linear distance derived from the tracking data. We fitted a separate LM for *A. femoralis* and *D. tinctorius* with sex and time period between recapture points as predictors. Contrary to the tracking data for *A. femoralis*, which was collected on an island with artificial pools, *A. femoralis* capture-recapture data were collected from a nearby natural population and thus served as additional validation for the effects of the island and artificial pools on the patterns of space use. We found that long-term movement extent distances based on capture-recapture data were larger in males than in females of *A. femoralis* (LM: $\beta_{\text{extent male}} = 0.55$, $t = 3.1$, $n = 165$, $P = 0.002$; Fig. S2a) and of *D. tinctorius* (LM: $\beta_{\text{extent male}} = 0.43$, $t = 2.62$, $n = 154$, $P = 0.01$; Fig S2b).

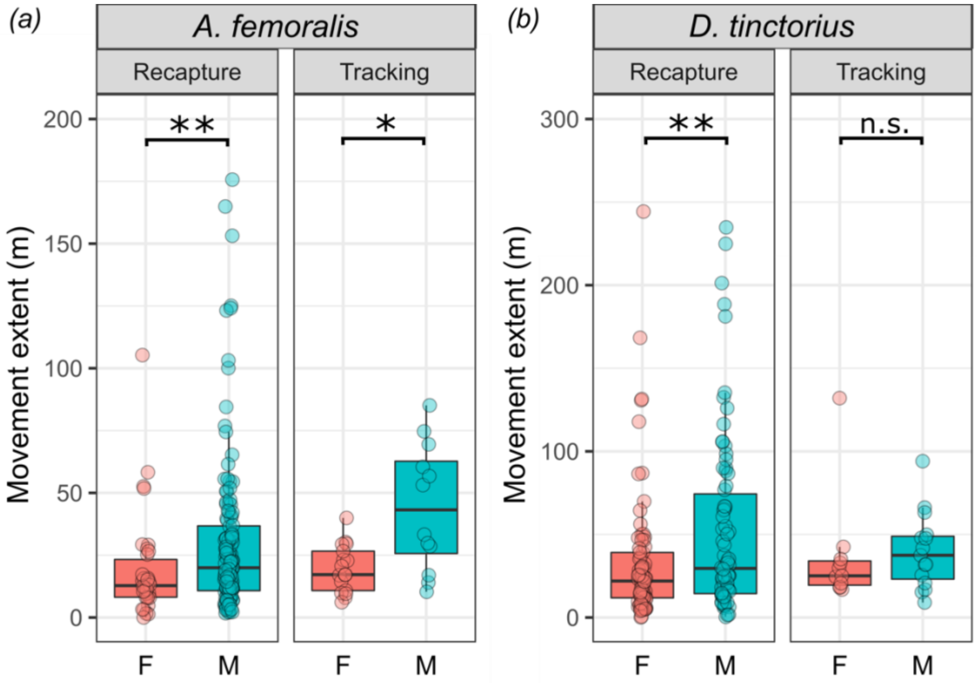


**Supplementary figure S2.** Comparison of sex differences in movement extent between long-term recapture data and short-term tracking. Boxplots showing sex differences in movement extent of (a) *A. femoralis* and (b) *D. tinctorius* female (F) and male (M) frogs calculated based on the long-term capture-recapture data (left panel) or short-term tracking data (right panel). Plot rectangles indicate the lower and upper quartiles (with the median as the dark line), whiskers extend to 1.5 times the interquartile range but do not extend past the full value range, and dots indicate individuals. Statistical significance levels are indicated as *p 0.05 – 0.01, **p <0.01, ***p <0.001.
