## Supplementary material for "Contrasting parental roles shape sex differences in poison frog space use but not navigational performance": Annex 2

### Navigational performance measured as angular deviation from the home direction

Trajectory straightness measures are influenced by the sampling rate, which was variable in our dataset. Therefore, we also quantified homing accuracy as an angular deviation from the home direction at fixed distance thresholds from the release site. The goal was to quantify the accuracy during the initial orientation near the release site, adjusted for translocation distance. We measured the angular deviation from the home center direction where the frog first crossed a 10 ∓5 meter or 40 ∓20 meter radius circle drawn around the release site for 50-m and 200-m translocation, respectively. This corresponded to ~ 20 % of the total translocation distance for each translocation distance. We compared the circular distributions of males and females with a MANOVA based on trigonometric functions (Landler et al., 2021) using the "manova" functions within the stats R package. We also evaluated if each group (males and females after 50-m or 200-m translocation) were significantly orientated towards home at the measured thresholds using the Rayleigh test of circular uniformity with the "rayleigh.test" function within the circular R package (Agostinelli & Lund, 2022). We found that after moving ~20 % of the translocation distance from the release site, *A. femoralis* males but not females were significantly oriented towards home (50-m: male Rayleigh $P < 0.001$, female Rayleigh $P = 0.15$; 200-m: male Rayleigh $P < 0.001$; Fig. S3a). Both males and females of *D. tinctorius* were significantly oriented when translocated 50-m (male Rayleigh $P = 0.007$, female Rayleigh $P < 0.001$; Fig. S3b), but not when translocated 200-m (male Rayleigh $P = 0.4$, female Rayleigh $P = 0.85$; Fig. S3e). Males but not females of *O. sylvatica* were significantly oriented when translocated 50-m (male Rayleigh $P = 0.015$, female Rayleigh $P = 0.18$; Fig. S3c), and *O. sylvatica* showed no homeward orientation after 200-m translocation (male Rayleigh $P = 0.9$, female Rayleigh $P = 0.9$; Fig. 7f). The only sex difference in the distribution of angular deviations revealed by MANOVA was in *A. femoralis* (50-m trans: $F(22) = 3.9$, $P = 0.03$).

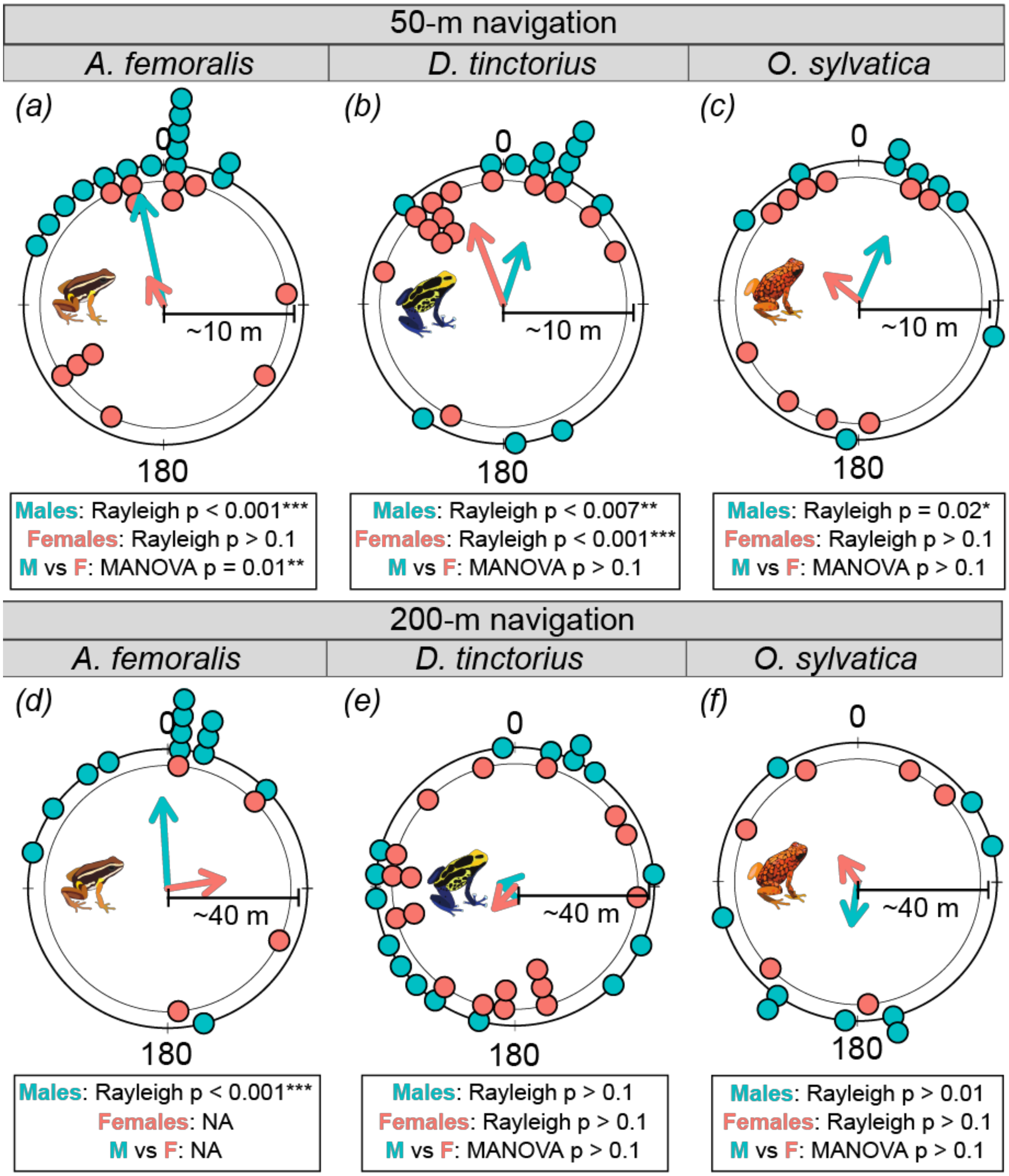


**Supplementary figure S3.** Sex differences in angular deviation from home direction measured at ~ 10 m or ~ 40 m from the release site for (a-c) 50-m and (d-f) 200-m translocation respectively. Dots on circular plots represent individual angular deviations with males (M) in teal and females (F) in red. Arrows represent the mean vector direction and length for each species, sex, and translocation distance. Circular statistics for significant deviation from uniformity are indicated separately for males and females together with a MANOVA directly comparing significant group differences. Statistical significance levels are indicated as *p 0.05 – 0.01, **p <0.01, ***p <0.001.
