## Supplementary material for "Contrasting parental roles shape sex differences in poison frog space use but not navigational performance": Annex 3

### Temporal patterns of homing

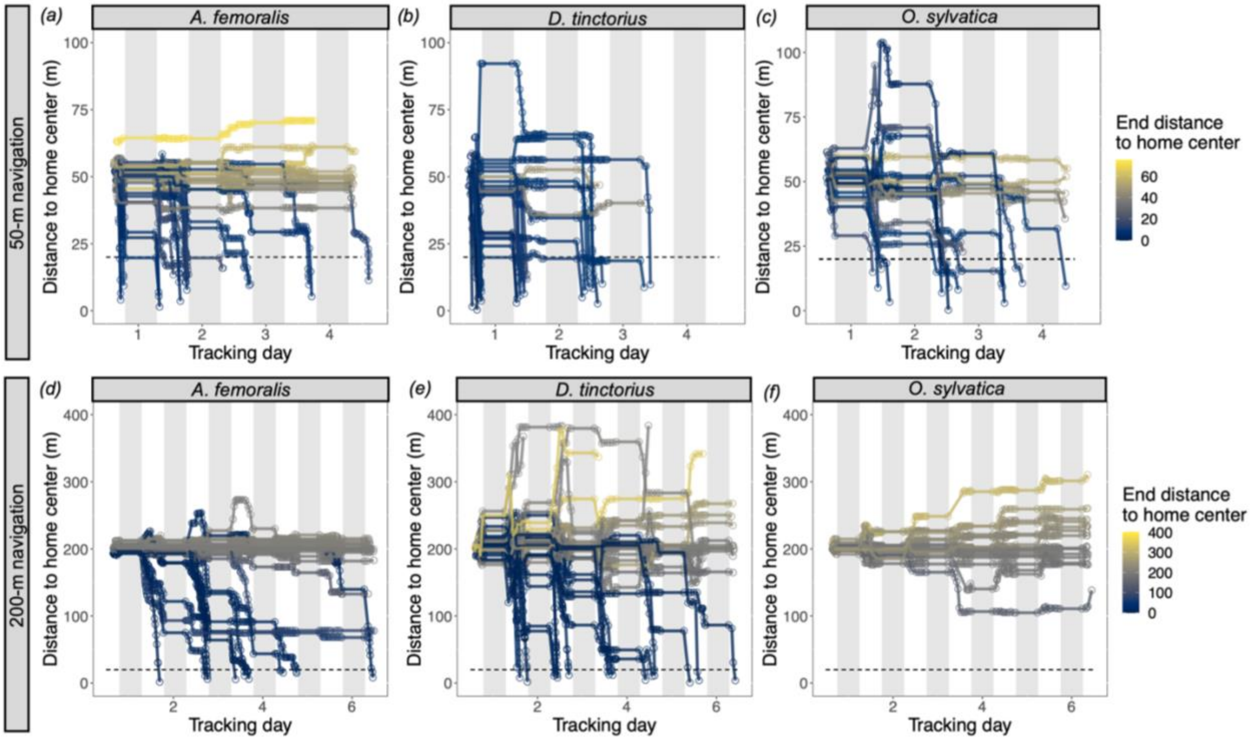


**Supplementary figure S4.** Species difference in temporal patterns of movement after (top row) 50-meter or (bottom row) 200-meter translocation. Y-axis shows distance (in meters) from the respective home center. X-axis shows time (in days) from the release after translocation. Color gradient indicates the distance remaining to the home area at the end of the tracking period. Blue represents the frogs that returned home, gray shows the frogs that stayed close to the release site or moved in parallel to home, and yellow shows frogs that moved away from home. Gray shaded areas represent night-time when frogs do not move. Each line represents a different individual. (a, d) *A. femoralis* (inconspicuous species) rarely moved unless moving towards home while (b, e) *D. tinctorius* and (c, f) *O. sylvatica* (two aposematic species) showed exploratory movements and often move away from home.
