## Supplementary material for "Contrasting parental roles shape sex differences in poison frog space use but not navigational performance": Annex 4

### Annex 4
### Supplementary statistics tables

**Supplementary table S1. Movement extent model summaries.**

| | *A. femoralis* $\log_e$ (movement extent) | | *D. tinctorius* $\log_e$ (movement extent) | | *O. sylvatica* $\log_e$ (movement extent) | |
|---|---|---|---|---|---|---|
| *Predictors* | *Estimates (CI)* | *p* | *Estimates (CI)* | *p* | *Estimates (CI)* | *p* |
| (Intercept) | 2.9 (0.65 – 5.2) | **0.014** | 5.9 (4.7 – 7.2) | **<0.001** | 3.4 (2.50 – 4.35) | **<0.001** |
| Sex [male] | 1.0 (0.2 – 1.8) | **0.019** | 0.6 (-0.45 – 1.7) | 0.240 | -0.8 (-1.3 – -0.4) | **0.001** |
| Tracking duration | 0.1 (-0.03 – 0.3) | 0.101 | -0.02 (-0.1 – 0.1) | 0.564 | 0.01 (-0.1 – 0.1) | 0.831 |
| Observations | 25 | | 19 | | 29 | |
| $R^2$ / $R^2$ adjusted | 0.34 / 0.28 | | 0.085 / -0.03 | | 0.38 / 0.33 | |

Statistical summary of three linear models with $\log_e$-transformed movement extent area in *A. femoralis*, *D. tinctorius*, and *O. sylvatica* as the response variable, sex as the predictor, and tracking duration (days) as a covariate. Statistical significance with $p < 0.05$ is highlighted in bold.

**Supplementary table S2. Explored area after 50-m translocation model summaries.**

| | *A. femoralis* 50-m $\log_e$ (explored area) | | *D. tinctorius* 50-m $\log_e$ (explored area) | | *O. sylvatica* 50-m $\log_e$ (explored area) | |
|---|---|---|---|---|---|---|
| *Predictors* | *Estimates* | *p* | *Estimates* | *p* | *Estimates* | *p* |
| (Intercept) | 17.5 (10.1 – 24.9) | **<0.001** | 5.7 (-0.5 – 11.8) | 0.068 | -19.5 (-84.4 – 45.5) | 0.530 |
| Sex [male] | 0.6 (0.3 – 0.9) | **<0.001** | 0.1 (-0.3 – 0.5) | 0.681 | 0.2 (-0.4 – 0.8) | 0.514 |
| Temp. | -0.5 (-0.8 – -0.3) | **<0.001** | 0.03 (-0.2 – 0.3) | 0.835 | 1.0 (-1.8 – 3.8) | 0.459 |
| Weight | 0.7 (-0.1 – 1.5) | 0.084 | 0.01 (-0.2 – 0.2) | 0.954 | 1.3 (-0.3 – 2.9) | 0.109 |
| Observations | 32 | | 31 | | 18 | |
| $R^2$ / $R^2$ adjusted | 0.59 / 0.55 | | 0.015 / -0.09 | | 0.23 / 0.06 | |

Statistical summary of three linear models with $\log_e$-transformed explored area in *A. femoralis*, *D. tinctorius*, and *O. sylvatica* as the response variable, sex as the predictor, and ambient daytime temperature (Temp.) and frog weight as covariates. Statistical significance with $p < 0.05$ is highlighted in bold.

**Supplementary table S3. Homing duration model summaries.**

| | *A. femoralis* 50-m log$_e$ (homing duration) | | *D. tinctorius* 50-m log$_e$ (homing duration) | | *D. tinctorius* 200-m log$_e$ (homing duration) | | *O. sylvatica* 50-m log$_e$ (homing duration) | |
|---|---|---|---|---|---|---|---|---|
| *Predictors* | *Estimates* | *p* | *Estimates* | *p* | *Estimates* | *p* | *Estimates* | *p* |
| (Intercept) | -23.0 (-47.2 – 1.2) | 0.061 | 2.3 (-15.95 – 20.5) | 0.801 | -2.8 (-19.7 – 14.1) | 0.720 | -70.8 (-140.2 – -1.45) | **0.046** |
| Sex [male] | -2.0 (-3.15 – -0.9) | **0.002** | -0.7 (-1.95 – 0.5) | 0.218 | -1.55 (-4.1 – 1.0) | 0.202 | 0.4 (-0.4 – 1.1) | 0.294 |
| Temp. | 1.2 (0.2 – 2.3) | **0.021** | 0.1 (-0.6 – 0.8) | 0.795 | 0.4 (-0.2 – 1.0) | 0.203 | 3.15 (0.15 – 6.1) | **0.042** |
| Weight | -1.7 (-4.3 – 0.85) | 0.175 | -0.6 (-1.2 – 0.05) | 0.069 | -0.5 (-1.8 – 0.7) | 0.364 | 0.2 (-2.05 – 2.4) | 0.860 |
| Observations | 19 | | 30 | | 14 | | 14 | |
| $R^2$ / $R^2$ adjusted | 0.54 / 0.44 | | 0.13 / 0.03 | | 0.31 / 0.10 | | 0.385 / 0.20 | |

Statistical summary of four linear models with log$_e$-transformed homing duration in *A. femoralis*, *D. tinctorius*, and *O. sylvatica* as the response variable, sex as the predictor, and ambient daytime temperature (Temp.) and frog weight as covariates. Statistical significance with $p < 0.05$ is highlighted in bold.

**Supplementary table S4. Sex difference in androgen levels.**

| | *A. femoralis* log$_e$ (androgens) | | *D. tinctorius* log$_e$ (androgens) | | *O. sylvatica* log$_e$ (androgens) | |
|---|---|---|---|---|---|---|
| *Predictors* | *Estimates* | *p* | *Estimates* | *p* | *Estimates* | *p* |
| (Intercept) | 2.42 (2.16 – 2.68) | **<0.001** | 3.16 (2.58 – 3.74) | **<0.001** | 2.26 (2.01 – 2.52) | **<0.001** |
| Sex [m] | 0.40 (0.09 – 0.72) | **0.012** | 1.04 (0.33 – 1.75) | **0.004** | 0.41 (0.08 – 0.74) | **0.015** |
| Time point [back home] | 0.12 (-0.19 – 0.44) | 0.444 | -0.16 (-0.71 – 0.38) | 0.556 | 0.07 (-0.21 – 0.35) | 0.631 |
| **Random Effects** | | | | | | |
| $\sigma^2$ | 0.42 | | 0.95 | | 0.24 | |
| $\tau_{00}$ | 0.00 $_{id}$ | | 0.52 $_{id}$ | | 0.08 $_{id}$ | |
| ICC | | | 0.35 | | 0.25 | |
| N | 35 $_{id}$ | | 35 $_{id}$ | | 34 $_{id}$ | |
| Observations | 66 | | 56 | | 51 | |
| Marginal $R^2$ / Conditional $R^2$ | 0.095 / NA | | 0.157 / 0.455 | | 0.125 / 0.342 | |

Statistical summary of three linear mixed models with log$_e$-transformed androgen levels in *A. femoralis*, *D. tinctorius*, and *O. sylvatica* as the response variable, sex and sampling point as the predictors, frog identity as the random factor. Statistical significance with $p < 0.05$ is highlighted in bold.

**Supplementary table S5. Androgen influence on spatial variables in *A. femoralis*.**

| | *A. femoralis* $\log_e$ (explored area) | | *A. femoralis* $\log_e$ (explored area) | | *A. femoralis* $\log_e$ (explored area) | |
|---|---|---|---|---|---|---|
| *Predictors* | *Estimates* | *p* | *Estimates* | *p* | *Estimates* | *p* |
| (Intercept) | 5.8 (5.4 – 6.1) | **<0.001** | 0.04 (0.02 – 0.1) | **0.014** | 0.9 (0.8 – 0.95) | **<0.001** |
| Androgens | 0.04 (-0.2 – 0.2) | 0.725 | 0.00 (-0.02 – 0.02) | 0.804 | 0.02 (-0.02 – 0.1) | 0.343 |
| Sex [male] | 0.3 (-0.2 – 0.8) | 0.261 | NA | | NA | |
| Translocation dist. [50m] | -0.7 (-1.1 – -0.3) | **0.002** | 0.03 (-0.02 – 0.01) | 0.259 | -0.1 (-0.2 – -0.03) | **0.029** |
| Homing success [1] | 1.2 (0.7 – 1.8) | **<0.001** | NA | | NA | |
| Temp. | NA | | -0.02 (-0.03 – -0.00) | 0.057 | -0.1 (-0.1 – -0.02) | **0.011** |
| Observations | 35 | | 12 | | 12 | |
| $R^2$ / $R^2$ adjusted | 0.67 / 0.62 | | 0.705 | | 0.61 | |

Statistical summary of three linear models with log-transformed explored area, homing duration, and homing trajectory straightness in *A. femoralis* as the response variables, androgen levels and sex as the predictors, translocation distance, homing success, and average daytime temperature (Temp.) as covariates. Statistical significance with $p < 0.05$ is highlighted in bold.

**Supplementary table S6. Androgen influence on spatial variables in *D. tinctorius.***

| | ***D. tinctorius* Log explored area** | | ***D. tinctorius* homing duration** | | ***D. tinctorius* trajectory straightness** | |
|---|---|---|---|---|---|---|
| *Predictors* | *Estimates* | *p* | *Estimates* | *p* | *Estimates* | *p* |
| (Intercept) | 7.24 (6.64 – 7.84) | **<0.001** | 0.02 (-0.02 – 0.09) | 0.462 | 0.52 (0.27 – 0.90) | **0.009** |
| Androgens | 0.33 (0.04 – 0.61) | **0.027** | 0.00 (-0.04 – 0.06) | 0.898 | -0.07 (-0.20 – 0.06) | 0.310 |
| Sex [male] | 0.92 (0.04 – 1.80) | **0.042** | 0.01 (-0.06 – 0.09) | 0.769 | -0.04 (-0.30 – 0.21) | 0.761 |
| Translocation dist. [50m] | -1.62 (-2.44 – -0.80) | **0.001** | 0.06 (-0.01 – 0.12) | 0.092 | 0.17 (-0.21 – 0.46) | 0.307 |
| Homing success [1] | 0.51 (-0.34 – 1.36) | 0.222 | NA | | NA | |

| | | | | | |
|---|---|---|---|---|---|
| Weight | 0.60<br>(0.18 – 1.02) | **0.008** | NA | | NA |
| Observations | 23 | | 16 | | 12 |
| $R^2$ / $R^2$ adjusted | 0.762 / 0.691 | | 0.308 | | 0.207 |

Statistical summary of three linear models with $\log_e$-transformed transformed explored area, homing duration, and homing trajectory straightness in *D. tinctorius* as the response variables, androgen levels and sex as the predictors, translocation distance, homing success, and frog weight as covariates. Statistical significance with p < 0.05 is highlighted in bold.

**Supplementary table S7. Androgen influence on spatial variables in *O. sylvatica*.**

| | *O. sylvatica*<br>Log explored area | | *O. sylvatica*<br>homing duration | | *O. sylvatica*<br>trajectory straightness | |
|---|---|---|---|---|---|---|
| *Predictors* | *Estimates* | *p* | *Estimates* | *p* | *Estimates* | *p* |
| (Intercept) | 6.40<br>(6.00 – 6.81) | **<0.001** | 0.05<br>(0.03 – 0.07) | **0.001** | 0.70<br>(0.54 – 0.91) | **<0.001** |
| Androgens | -0.06<br>(-0.31 – 0.18) | 0.589 | 0.01<br>(-0.00 – 0.02) | 0.188 | 0.21<br>(0.09 – 0.32) | **0.003** |
| Sex [male] | 0.23<br>(-0.25 – 0.71) | 0.333 | -0.01<br>(-0.03 – 0.02) | 0.549 | -0.13<br>(-0.35 – 0.09) | 0.219 |
| Translocation dist. [50m] | -0.99<br>(-1.62 – -0.37) | **0.003** | NA | | NA | |
| Homing success [1] | 1.07<br>(0.47 – 1.67) | **0.001** | NA | | NA | |
| Observations | 28 | | 13 | | 13 | |
| $R^2$ / $R^2$ adjusted | 0.433 / 0.334 | | 0.113 | | 0.451 | |

Statistical summary of three linear models with $\log_e$-transformed transformed explored area, homing duration, and homing trajectory straightness in *O. sylvatica* as the response variables, androgen levels and sex as the predictors, translocation distance and homing success as covariates. Statistical significance with p < 0.05 is highlighted in bold.

**Supplementary table S8. Exploration influence on delta androgen levels.**

| | *A. femoralis*<br>delta androgens | | *D. tinctorius*<br>delta androgens | | *O. sylvatica*<br>delta androgens | |
|---|---|---|---|---|---|---|
| *Predictors* | *Estimates* | *p* | *Estimates* | *p* | *Estimates* | *p* |
| (Intercept) | 7.84<br>(-2.75 – 18.44) | 0.140 | -47.48<br>(-382.53 – 287.57) | 0.768 | 2.64<br>(-12.97 – 18.25) | 0.717 |
| Explored area | 12.70<br>(3.37 – 22.03) | **0.010** | 12.77<br>(-107.77 – 133.31) | 0.825 | 0.53<br>(-7.61 – 8.67) | 0.889 |
| Sex [male] | 12.70<br>(-3.73 – 29.13) | 0.124 | 41.74<br>(-144.22 – 227.70) | 0.641 | -6.99<br>(-21.00 – 7.03) | 0.296 |

| | | | | | | |
|---|---|---|---|---|---|---|
| Tracking duration | -7.18 (-14.87 – 0.52) | 0.066 | 4.14 (-187.48 – 195.76) | 0.964 | 4.01 (-10.97 – 18.99) | 0.567 |
| Homing success [1] | -28.63 (-53.37 – -3.88) | **0.025** | 39.40 (-449.42 – 528.21) | 0.866 | 6.24 (-22.51 – 34.99) | 0.642 |
| Observations | 31 | | 21 | | 16 | |
| $R^2$ / $R^2$ adjusted | 0.353 / 0.254 | | 0.027 / -0.216 | | 0.141 / -0.171 | |

Statistical summary of three linear models with delta androgen levels in *A. femoralis*, *D. tinctorius*, and *O. sylvatica* as the response variable, explored area and sex as the predictors, tracking duration and homing success as covariates. Statistical significance with $p < 0.05$ is highlighted in bold.
