## Supplementary material for "Contrasting parental roles shape sex differences in poison frog space use but not navigational performance": Annex 5

### Frog tagging and tracking methods

We used two previously described tracking methods: harmonic direction-finding (HDF) with passive transponders for *A. femoralis* and *O. sylvatica* (for more details see Beck et al., 2017; Fischer et al., 2020; Pašukonis et al., 2014) and radio-tracking with Very High Frequency (VHF) transmitters for the larger *D. tinctorius* (for more details see Pašukonis et al., 2018, 2019). For HDF, we used a handheld transceiver (R8 and R9, Recco AB, Lindigö, Sweden) to detect miniature transponders attached to frogs with a silicone waistband. We used custom-made transponders with a thin flexible ~ 10 - 14 cm wire antenna dragging freely behind the frog as well as commercial animal reflectors (R-30CL, Recco AB, Lindigö, Sweden) that consist of a diode encased in a thin, flexible 3x65 mm film. The tags with waistbands constituted ~ 5% of adult *A. femoralis* and *O. sylvatica* body weight (tag weight ~ 0.1 g; *A. femoralis* female median weight = 2 g, n = 32; male median weight 1.8 g, n = 30; *O. sylvatica* la Florida median weight = 1.8 g, n = 30, no sex difference; Canandé median weight = 2.2 g, n = 51, no sex difference). *Dendrobates tinctorius* were equipped with miniature VHF transmitters (BD2X, Holohil Systems Ltd., Carp, ON, Canada; NTQ2, Lotek Wireless, Newmarket, ON, Canada; PicoPip, Biotrack, Wareham, UK; V5, Telemetrie-Service Dessau, Dessau-Rosslau, Germany). Frogs were located using a portable radio-tracking receiver (Sika, Biotrack Ltd.) and a flexible Yagi-antenna (Biotrack Ltd.). The tags constituted 6 – 12 % of adult *D. tinctorius* body weight (tag weight 0.35 – 0.4 g; female median weight = 5.5 g, n = 44; male median weight = 3.9 g, n = 37).

We tagged and tracked a total of 311 frogs. Frogs were captured, photographed, measured, tagged, and released at the capture locations within 30 min, with a few longer exceptions of up to 2 hours. Handling itself usually took 5 to 15 min and frogs were kept in a plastic bag or net cage after capture until an experimenter was available for tagging. After release, we located each frog multiple times a day, recorded their position, and observed behavior. We tried to visually spot the frog with every measurement, although this was not always possible due to dense habitat and, in these cases, we narrowed the location to approximately one meter. To map frog movements, we measured distance and direction from a fixed reference point using laser distance meters (GLM 40, Bosch, Grelingen, Germany and DW0165, DeWalt, Towson, USA) and precision compasses (Suunto KB20 and Suunto Tandem, Suunto, Vantaa, Finland). Frog locations were recorded on a digital GIS map of the study area using handheld GIS/GPS devices (Vanquisher SV-86 rugged Windows tablet, Sinicvision Technology Co., Shenzhen, China; WinTab 9, Odys, Willich, Germany; MobileMapper10; SpectraPrecision, Westminster, CO, USA) and GIS software ArcPad 10 (ESRI, Redlands, USA). Occasionally, when frogs moved out of the mapped area, we recorded their location by averaging at least 30 GPS points with a GPS/GIS device MobileMapper 10. All data were collected as GIS spatial points with associated behavioral information. These data were sorted, visualized, and error-checked in GIS software QGIS (versions 2.14 and 3.18, QGIS.org, 2022) and ArcMap (various versions, ESRI, Redlands, USA). All data were checked point-by-point at least twice and corrected for errors such as wrong frog identities, duplicates, and impossible locations. All suspect points where frog identity or location could not be unambiguously confirmed and corrected were removed.

Tagged frogs showed full mobility and the full range of natural behaviors, but tagging-related issues occasionally required experimenter intervention. Antennas occasionally tangled and snagged on the vegetation, in which case we intervened to release the frog. This was especially the case in *O. sylvatica*, which often climbed on vegetation. Frogs were shortly handled every few days to check the tag fit, and the tag was adjusted or removed as necessary. Five frogs died during the study: one due to experimenter error (trampling), one due to the tag (snagged antenna on the vegetation), two due to predation, and one for unknown reasons. Some frogs (16 %) experienced skin damage ranging from superficial abrasions to deep open wounds from attachment. The prevalence of wounds varied between species (*A. femoralis* 6 %, *O. sylvatica* 18 %, *D. tinctorius* 25 %), and field seasons. Fast recovery of skin injuries was observed in the frogs that were recaptured after removing the tag. To evaluate if skin injuries affected frog mobility, we categorized the injuries into two categories of severity, superficial skin abrasion and skin lesion. We then included injury as a categorical predictor in models of home range size, movement extent, daily travel, explored area, and homing duration. In space use models, the injury had a significant effect on home range size and daily travel in *A. femoralis*, but not in *D. tinctorius* and *O. sylvatica*. In the navigation models, the injury had a significant effect in the model of explored area size in *O. sylvatica* translocated 200 meters. Based on these results, we excluded *A. femoralis* (1 male and 1 female) and *O. sylvatica* (3 males and 3 females) with skin lesions from space use and navigation analyses, respectively. The overall results of all models remained the same with and without injured frogs, but we report the conservative values after excluding these individuals.
