## Supplementary material for "Contrasting parental roles shape sex differences in poison frog space use but not navigational performance": Annex 6

### Annex 6
### Supplementary methods tables

**Supplementary table S9. Study site characteristics.**

| Site name | Locality | Coordinates | Area (ha) | Site characteristic | Breeding sites |
|---|---|---|---|---|---|
| Nouragues island | Nouragues Nature Reserve, French Guiana | 4°02' N, 52°41' W | 4.6 | Free-range experimental island population | Mostly artificial |
| Nouragues main | Nouragues Nature Reserve, French Guiana | 4°02' N, 52°41' W | 25 | Natural population | Mostly natural |
| Florida | La Florida, Ecuador | 0°15' S, 79°02' W | 0.5 | Free-range experimental enclosure population | Natural and artificial |
| Canandé | Reserva Canandé, Ecuador | 0°32' N, 79°13' W | 5.5 | Natural population | Natural |

**Supplementary table S10. Data type and dataset descriptive.**

| Data types | Dataset id | Study period | Species | Site | N frogs tagged | N frogs included | Note |
|---|---|---|---|---|---|---|---|
| Space use tracking | af16 | 2016/02/08 - 2016/03/20 | *A. femoralis* | Nouragues island | 24 | 17 | Only females tracked |
| | af18 | 2018/03/17 - 2018/04/18 | *A. femoralis* | Nouragues island | 12 | 12 | Only males tracked |
| | dt16 | 2016/01/27 - 2016/03/19 | *D. tinctorius* | Nouragues main | 31 | 26 | |
| | os17 | 2017/04/23 - 2017/05/16 | *O. sylvatica* | Florida | 31 | 29 | |
| Space use recapture | dt_recap | 2009/01/09 - 2011/06/05 | *D. tinctorius* | Nouragues main | *NA* | 154 | Long-term recapture data |
| | af_recap | 2014/01/24 - 2019/04/22 | *A. femoralis* | Nouragues main | *NA* | 165 | Long-term recapture data |
| Navigation | af17 | 2017/01/27 - 2017/03/11 | *A. femoralis* | Nouragues main | 34 | 28 | |
| | dt17 | 2017/02/26 - 2017/03/23 | *D. tinctorius* | Nouragues main | 36 | 29 | |
| Navigation, androgens, space use tracking | os19 | 2019/05/21 - 2019/06/25 | *O. sylvatica* | Canandé | 52 | 39 | Dataset also used for space use validation |
| Navigation, androgens | dt19 | 2019/02/19 - 2019/03/26 | *D. tinctorius* | Nouragues main | 47 | 38 | |
| | af20 | 2020/01/21 - 2020/03/16 | *A. femoralis* | Nouragues main | 44 | 36 | |

**Supplementary table S11. Dependent variables definitions used in the statistical analyses.**

| Data types | Variable name | Definition | Datasets |
|---|---|---|---|
| Space use | home range | Area under the 95% utilization density contour | af16, af18, dt16, os17 (tracked > 7 days) |
| | extent area | Area under the minimum convex polygon | af16, af18, dt16, os17 (tracked > 7 days) |
| | daily travel | Cumulative daily distance traveled | af16, af18, dt16, os17, os19 (tracked > 2 days) |
| | extent distance | Maximum linear distance between locations of one individual | af16, af18, dt16, os17, os19 (tracked > 2 days) dt_recap, af_recap (recaptured >30 days) |
| Navigation androgens | homing success | Homing (yes/no) at least 70% of translocation distance | af17, dt17, os19, dt19, af20 (all translocated frogs) |
| | explored area | Area within five meters of the movement trajectory | af17, dt17, os19, dt19, af20 (all translocated frogs) |
| | trajectory straightness | Ratio between straight-line and cumulative distance from the release site to the end of the homing trajectory | af17, dt17, os19, dt19, af20 (successfully homing frogs) |
| | homing duration | Daytime hours from the release time to the arrival within 10-meters of the home area polygon | af17, dt17, os19, dt19, af20 (homing frogs) |
| Navigation | angular deviation | Angular deviation from home center direction measured at ~20% of the translocation distance from the release site | af17, dt17, os19, dt19, af20 (all translocated frogs) |
| Androgens | androgen level | Water-borne androgen concentration measured within 2 day before translocation and after returning home | os19, dt19, af20 (all translocated frogs) |
| | delta androgens | Difference between androgen concentrations measured back at the home site and baseline androgens. | os19, dt19, af20 (all translocated frogs) |

**Supplementary table S12. Predictor variable definitions use in the statistical analyses.**

| Predictor | Definition |
|---|---|
| Species | Three level factor: *A. femoralis, D. tinctorius, O. sylvatica* |
| Sex | Two level factor: male, female |
| Behavior | Three or two level factor: parental, mating, other |
| Weight | Continuous numeric: frog weight in grams |
| Daytime temperature | Continuous numeric: average daytime temperature measured between sunrise and sunset |
| Mean temperature | Continuous numeric: daytime temperature averaged over the entire tracking period of an individual |
| Translocation distance | Two level factor: 50-m, 200-m |
| Homing success | Two level factor: yes, no |
| Explored area | Continuous numeric: area within five meters of the movement trajectory |
| Tracking duration | Continuous numeric: total tracking duration of an individual |
| Period | Count numeric: Number of days between recaptures of the same individual |
| Baseline androgens | Continuous numeric: Water-borne androgen concentration measured within 2 day before translocation |
| Time point | Two level factor: baseline, back home |

**Supplementary table S13. All models performed for the statistical analyses.**

| Data type | Dependent variable | Predictors | Model id | Datasets | Data subset |
|---|---|---|---|---|---|
| Space use | Log$_e$ (home range) | species, sex, tracking duration | m11_allud | af16 + af18 + dt16 + os17 | All species |
| | | sex, tracking duration | m12_afud | af16 + af18 | *A. femoralis* |
| | | sex, tracking duration | m13_dtud | dt16 | *D. tinctorius* |
| | | sex, tracking duration | m14_osud | os17 | *O. sylvatica* |
| | Log$_e$ (extent area) | species, sex, tracking duration | m15_allmcp | af16 + af18 + dt16 + os17 | All species |
| | | sex, tracking duration | m16_afmcp | af16 + af18 | *A. femoralis* |
| | | sex, tracking duration | m17_dtmcp | dt16 | *D. tinctorius* |
| | | sex, tracking duration | m18_osmcp | os17 | *O. sylvatica* |
| | Log$_e$ (daytime travel) | sex, species, points per day | m1_alldl | af16 + af18 + dt16 + os17 | All species |
| | | sex, behavior, daytime temperature, points per day[1] | m2_afdl | af16 + af18 | *A. femoralis* |
| | | behavior, daytime temperature | m4_affdl | af16 | *A. femoralis* females |
| | | behavior, daytime temperature | m3_afmdl | af18 | *A. femoralis* males |
| | | sex, behavior, daytime temperature, points per day[1] | m5_dtdl | dt16 | *D. tinctorius* |
| | | behavior[2] | m7_dtfl | dt16 | *D. tinctorius* females |
| | | behavior[2] | m6_dtml | dt16 | *D. tinctorius* males |
| | | sex, behavior, daytime temperature, points per day[1] | m8_osdl | os17 | *O. sylvatica* |
| | | behavior[2] | m10_osmdl | os17 | *O. sylvatica* females |
| | | behavior[2] | m9_osfdl | os17 | *O. sylvatica* males |
| Space use validation | Log$_e$ (extent distance) | sex, tracking duration | m19_aftr | af16 + af18 | *A. femoralis* tracking |
| | | sex, period | m20_afcr | af_recap | *A. femoralis* recapture |
| | | sex, tracking duration | m21_dttr | dt16 | *D. tinctorius* tracking |
| | | sex, period | m22_dtcr | dt_recap | *D. tinctorius* recapture |
| | | sex, tracking duration | m23_oscan | os19 | *O. sylvatica* tracking natural site |
| | | sex, tracking duration | m24_osoto | os17 | *O. sylvatica* tracking enclosures |
| | Log$_e$ (daytime travel) | sex | m25_oscan | os19 | *O. sylvatica* tracking natural site |
| | | sex | m26_osoto | os17 | *O. sylvatica* tracking enclosures |
| Navigation | Homing success | sex, weight, mean temperature | m1_afprob | af17 + af20 | *A. femoralis* 50-m |
| | | sex, weight, mean temperature | m2_dtprob | dt17 + dt19 | *D. tinctorius* 200-m |
| | | sex, mean temperature[3] | m3_osprob | os19 | *O. sylvatica* 50-m |
| | Log$_e$ | sex, weight, mean temperature | m4_af50ex | af17 + af20 | *A. femoralis* 50-m |

| | | | | | |
|---|---|---|---|---|---|
| | (explored area) | sex, weight, mean temperature | m5_af200ex | af17 + af20 | *A. femoralis* 200-m |
| | | sex, weight, mean temperature | m6_dt50ex | dt17 + dt19 | *D. tinctorius* 50-m |
| | | sex, weight, mean temperature | m7_dt200ex | dt17 + dt19 | *D. tinctorius* 200-m |
| | | sex, weight, mean temperature | m8_os50ex | os19 | *O. sylvatica* 50-m |
| | | sex, weight, mean temperature | m9_os200ex | os19 | *O. sylvatica* 200-m |
| | Trajectory straightness | sex, mean temperature | m10_af50sc | af17 + af20 | *A. femoralis* 50-m homing |
| | | sex, mean temperature | m11_dt50sc | dt17 + dt19 | *D. tinctorius* 50-m homing |
| | | sex, mean temperature | m12_dt200sc | dt17 + dt19 | *D. tinctorius* 200-m homing |
| | | sex, mean temperature | m13_os50sc | os19 | *O. sylvatica* 50-m homing |
| | $Log_e$ (homing duration) | sex, weight, mean temperature | m14_af50dr | af17 + af20 | *A. femoralis* 50-m homing |
| | | sex, weight, mean temperature | m15_dt50dr | dt17 + dt19 | *D. tinctorius* 50-m homing |
| | | sex, weight, mean temperature | m16_dt200dr | dt17 + dt19 | *D. tinctorius* 200-m homing |
| | | sex, weight, mean temperature | m17_os50dr | os19 | *O. sylvatica* 50-m homing |
| | Angular deviation | sex | m18_af50cr | af17 + af20 | *A. femoralis* 50-m |
| | | sex | m19_dt50cr | dt17 + dt19 | *D. tinctorius* 50-m |
| | | sex | m20_dt200cr | dt17 + dt19 | *D. tinctorius* 200-m |
| | | sex | m21_os50cr | os19 | *O. sylvatica* 50-m |
| Androgens | $Log_e$ (baseline androgens) | sex, time point | m1_afT | af20 | *A. femoralis* |
| | | sex, time point | m1_dtT | dt19 | *D. tinctorius* |
| | | sex, time point | m1_osT | os19 | *O. sylvatica* |
| | Homing success | baseline androgens, sex, translocation distance[4] | m2_afT | af20 | *A. femoralis* |
| | | baseline androgens, sex, translocation distance[4] | m2_dtT | dt19 | *D. tinctorius* |
| | | baseline androgens, sex, translocation distance, weight[4] | m1_osT | os19 | *O. sylvatica* |
| | $Log_e$ (explored area) | baseline androgens, sex, translocation distance, homing success[4] | m3_afT | af20 | *A. femoralis* |
| | | baseline androgens, sex, translocation distance, homing success, weight[4] | m3_dtT | dt19 | *D. tinctorius* |
| | | baseline androgens, sex, translocation distance, homing success[4] | m3_osT | os19 | *O. sylvatica* |
| | Homing duration | baseline androgens, translocation distance, mean temperature[4] | m4_afT | af20 | *A. femoralis* homing |
| | | baseline androgens, sex, translocation distance[4] | m4_dtT | dt19 | *D. tinctorius* homing |
| | | baseline androgens, sex[4] | m4_osT | os19 | *O. sylvatica* homing |
| | Trajectory straightness | baseline androgens, translocation distance, mean temperature[4] | m5_afT | af20 | *A. femoralis* homing |

| | | | | | |
|---|---|---|---|---|---|
| | | baseline androgens, sex, translocation distance[4] | m5_dtT | dt19 | *D. tinctorius* homing |
| | | baseline androgens, sex[4] | m5_osT | os19 | *O. sylvatica* homing |
| | Delta androgens | explored area, sex, tracking duration, homing success[4] | m6_afT | af20 | *A. femoralis* homing |
| | | explored area, sex, tracking duration, homing success[4] | m6_dtT | dt19 | *D. tinctorius* homing |
| | | explored area, sex, tracking duration, homing success[4] | m6_osT | os19 | *O. sylvatica* homing |

[1] Full model.

[2] Daytime temperature excluded based on model selection for both sexes.

[3] Weight excluded because the model with weight didn't converge.

[4] Weight and/or mean temperature were excluded to reduce the number of predictors if the model excluding these predictors (separately) was not significantly different ($P > 0.1$) from the full model.
